# Interplay between PML NBs and HIRA for H3.3 dynamics following type I interferon stimulus

**DOI:** 10.1101/2021.11.30.470516

**Authors:** Constance Kleijwegt, Florent Bressac, Camille Cohen, Pascale Texier, Thomas Simonet, Laurent Schaeffer, Patrick Lomonte, Armelle Corpet

## Abstract

Promyelocytic Leukemia Nuclear Bodies (PML NBs) are nuclear membrane-less organelles physically associated with chromatin underscoring their crucial role in genome function. The H3.3 histone chaperone complex HIRA accumulates in PML NBs upon senescence, viral infection or IFN-I treatment in primary cells. Yet, the molecular mechanisms of this partitioning and its function in regulating histone dynamics have remained elusive. By using specific approaches, we identify intermolecular SUMO-SIM interactions as an essential mechanism for HIRA recruitment in PML NBs. Upon IFN-I stimulation, PML is required for interferon-stimulated genes (ISGs) transcription and PML NBs become juxtaposed to ISGs loci at late time points of IFN-I treatment. HIRA and PML are necessary for the prolonged H3.3 deposition at the transcriptional end sites of ISGs, well beyond the peak of transcription. Though, HIRA accumulation in PML NBs is dispensable for H3.3 deposition on ISGs. Hence, we describe an independent role of PML NBs as nuclear depot centers to regulate HIRA distribution in the nucleus, as a consequence of the availability of the pool of H3.3-H4 dimers or chromatin compaction. We thus uncover a dual function for PML/PML NBs in regulating ISGs transcription and H3.3 deposition at ISGs, and in modulating the nuclear distribution of HIRA upon inflammatory response.

## Introduction

Promyelocytic Leukemia Nuclear Bodies (PML NBs) are membrane-less organelles, also called biomolecular condensates (Banani et al. 2017), that concentrate proteins at discrete sites within the nucleoplasm thus participating in the spatio-temporal control of biochemical reactions (Lallemand-Breitenbach and de Thé 2018; Corpet et al. 2020; Li et al. 2020). PML NBs are 0.1-1μm diameter hollow sphere structures that vary in size and number depending on cell type, cell-cycle phase, or physiological state, highlighting their stress-responsive nature. The tumor-suppressor PML protein is the primary scaffold of PML NBs and forms an outer shell, together with the SP100 nuclear antigen, surrounding an inner core of dozens of proteins that localize constitutively or transiently in PML NBs. PML (also known as TRIM19) is a member of the tripartite motif (TRIM)-containing protein superfamily characterized by a conserved N-terminal RBCC motif essential for PML polymerization. Several isoforms of PML exist, all containing the RBCC motif and three well-characterized small-ubiquitin-related modifier (SUMO) modification sites at lysines K65, K160 and K490 and a SUMO interacting motif (SIM) enabling its interaction with SUMOylated proteins (Uggè et al. 2022; Corpet et al. 2020). SUMO E2 conjugating enzyme UBC9-mediated SUMOylation of PML enforces PML-PML interactions via intermolecular SUMO-SIM interactions. It also drives the multivalent recruitment of inner core protein clients through their SIM, possibly via liquid-liquid phase separation (LLPS) mechanisms (Corpet et al. 2020; Li et al. 2020; Sahin et al. 2014).

PML NBs have been involved in a wide variety of biological processes such as senescence, antiviral response, DNA damage response, or stemness suggesting that they are fully significant structures. The molecular mechanisms through which they exert their broad physiological impact are not fully elucidated yet. While PML NBs are in general devoid of DNA, except in specific cases (for review (Corpet et al. 2020)), they reside in the interchromatin nuclear space (Boisvert et al. 2000) and can associate with specific genomic loci (Shiels et al. 2001; Wang 2004; Kurihara et al. 2020; Ching et al. 2013; Kumar et al. 2007; Chang et al. 2013; Delbarre et al. 2017). PML NBs have been found associated with both transcriptionally-active domains (Boisvert et al. 2000; Wang 2004; Kurihara et al. 2020), as well as heterochromatin regions such as telomeres suggesting an important function in chromatin domain organization and regulation of their transcriptional state (for review (Delbarre and Janicki 2021)).

Targeted deposition of histones variants is crucial for chromatin homeostasis and the maintenance of cell identity (Allis and Jenuwein 2016). Among histone H3 variants, H3.3 is expressed throughout the cell cycle and is incorporated onto DNA in a DNA-synthesis independent manner by dedicated histone chaperone complexes (Martire and Banaszynski 2020). Histone cell cycle regulator A (HIRA) chaperone complex, composed of HIRA, ubinuclein 1 or ubinuclein 2 (UBN1 or UBN2) and calcineurin-binding protein CABIN1 is responsible for H3.3 deposition in transcriptionally active regions including enhancers, promoters and gene bodies, as well as in nucleosome-free regions and DNA damage sites (Ray-Gallet et al. 2002; 2011; Goldberg et al. 2010; Zhang et al. 2017) (for review (Martire and Banaszynski 2020; Ricketts and Marmorstein 2016)). HIRA has also been recently shown to be involved in the transcription-mediated recycling of parental H3.3 (Torné et al. 2020). While HIRA complex is diffusively distributed in the nuclei of proliferating somatic cells, it relocalizes in PML NBs upon various stresses such as senescence entry (Rai et al. 2011; Zhang et al. 2005; Banumathy et al. 2009; Jiang et al. 2011), viral infection (Cohen et al. 2018; McFarlane et al. 2019; Rai et al. 2017), or interferon type I (IFN-I) treatment (Rai et al. 2017; McFarlane et al. 2019). These latter events underscore a role of HIRA in intrinsic anti-viral defense via chromatinization of incoming viral genomes (Rai et al. 2017; Cohen et al. 2018) as well as stimulation of innate immune defenses in the case of viral infection (McFarlane et al. 2019).

However, the exact significance of HIRA localization in PML NBs upon inflammatory stress response, as well as the role of the PML NBs themselves, remain to be defined. PML NBs may act as buffering/sequestration/degradation structures for various chromatin-related proteins, and be a means to target them to specific chromatin regions juxtaposed to PML NBs. Here we investigated the molecular mechanisms of HIRA localization in PML NBs. We show that HIRA localizes in PML NBs in a SIM-SUMO-dependent manner upon IFN-I treatment. We provide evidence that PML is required for interferon-stimulated genes (ISGs) expression, and that ISGs loci juxtapose to PML NBs. ChIP-Seq analysis reveals a long-lasting H3.3 deposition on the 3’ end of ISGs, which is partly dependent on HIRA and PML, but independent of HIRA localization in PML NBs. Instead, we uncover that PML NBs rather act as nuclear depot centers for HIRA, depending upon histone availability and dynamics. Together, our results put forward a dual role for PML/PML NBs during the inflammatory response both regulating ISGs transcription as well as H3.3 deposition at ISGs, and acting as storage centers to modulate HIRA availability on chromatin.

## Results

### HIRA accumulation in PML NBs correlates with an increase in PML valency

Several stimuli, such as IFN-I treatment (Rai et al. 2017; McFarlane et al. 2019), can trigger HIRA accumulation in PML NBs. Previous studies suggest that a valency-dependent switch-like partitioning of a given client protein in the condensed PML NB phase is controlled by the multivalent interactions between its SIM motifs and SUMOylated lysines on the PML protein, as shown for the H3.3 histone chaperone DAXX (Banani et al. 2016; Sahin et al. 2014), which localizes constitutively in PML NBs (Ishov et al. 1999). As a first step towards deciphering the mechanism of HIRA localization in PML NBs, we investigated the valency-dependent recruitment of HIRA in PML NBs. Treatment of human primary foreskin diploid fibroblast BJ cells with the TLR3 ligand poly(I:C), a strong stimulant of the IFN-I pathway, or with the Tumor necrosis factor α (TNFα) cytokine, triggered a strong accumulation of HIRA in PML NBs (Figures 1A-B), similarly to a control IFN-I treatment (Sup. Figure 1A). The accumulation was abrogated by addition of ruxolitinib, an inhibitor of the JAK-STAT pathway downstream of the IFN-I receptor, underscoring the involvement of the IFN-I signaling pathway in primary cells (Figures 1A-B and Sup. Figure 1A). IFNβ, poly(I:C) and TNFα induced an IFN-I dependent increase of PML and its SUMOylated forms (Figure 1C), confirming previous data (Stadler et al. 1995; Gao et al. 2008). Treatment with other pro-inflammatory cytokines such as IL-6 or the IL-8 chemokine, did not increase PML protein levels or SUMOylation (Sup. Figure 1B), nor affected HIRA localization that remained pan-nuclear (Sup. Figure 1C). These results suggest that an IFN-I-dependent increase of PML valency (increase in protein levels and SUMOylation), but not of HIRA (Sup. Figure 1D) is part of the mechanism for HIRA accumulation in PML NBs.

**Figure 1.**
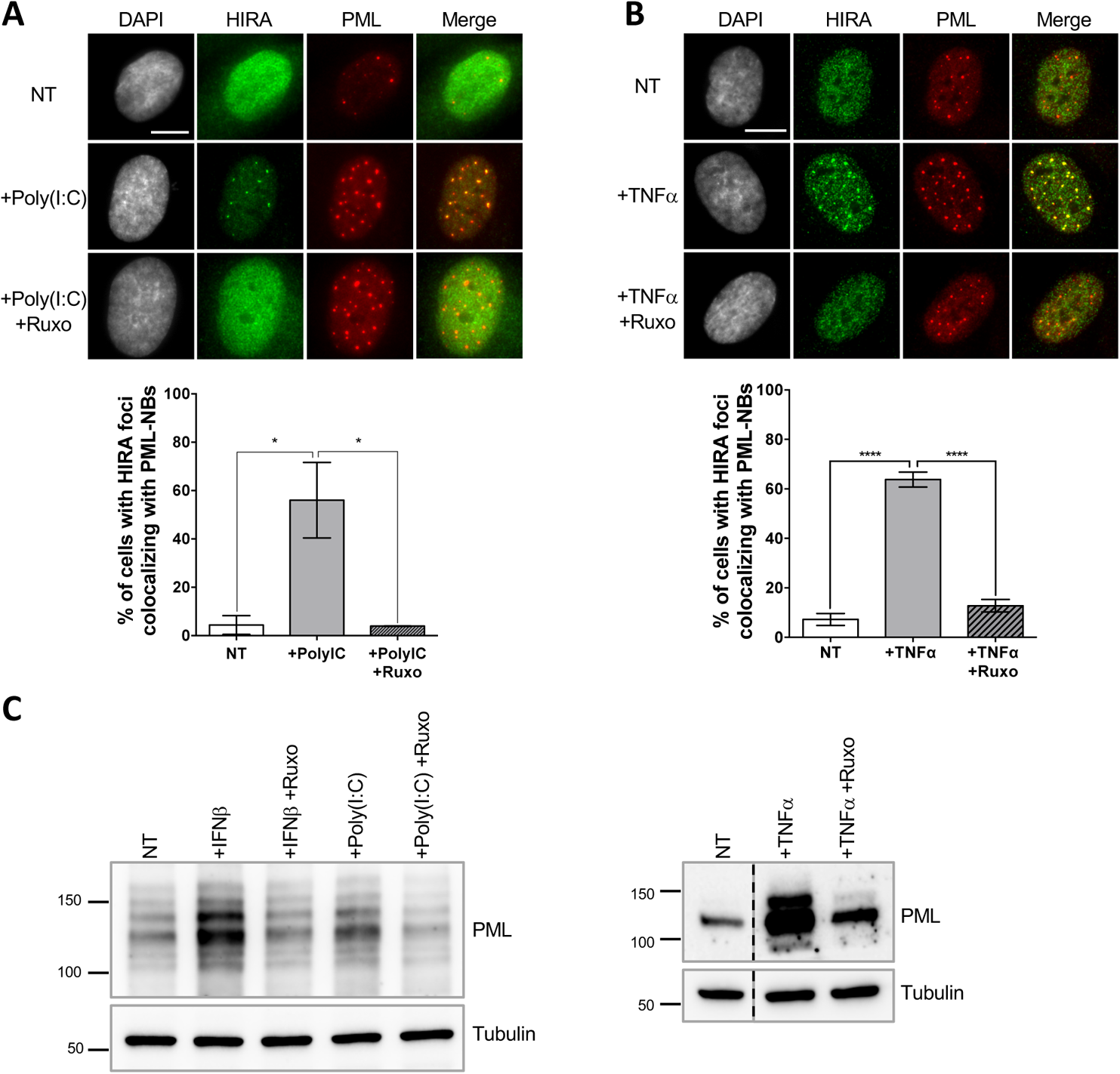
HIRA accumulation in PML NBs correlates with increased PML valency in primary cells. **A-B**. (top) Fluorescence microscopy visualization of HIRA (green) and PML (red) in BJ cells treated with Poly(I:C) at 10μg/mL for 24h (left) or with TNFα at 100ng/mL for 24h (right). Ruxolitinib (Ruxo) was added at 2μM one hour before Poly(I:C) or TNFα treatment and for 25h. Cell nuclei are visualized by DAPI staining (grey). Scale bars represent 10μm. (bottom) Histograms show quantitative analysis of cells with HIRA localization at PML NBs. p-values (Student t-test): *<0,05; ****<0,0001. Numbers on all histograms represent the mean of 3 independent experiments (±SD). **C.** (left) Western blot visualization of PML from total cell extracts of BJ cells treated with IFNβ at 1000U/mL or Poly(I:C) for 24h and with ruxolitinib (Ruxo) at 2µM one hour before treatment and for 25h. (right) Western blot visualization of PML from RIPA extracts of BJ cells treated with TNFα at 100ng/mL for 24h and with ruxolitinib (Ruxo) at 2µM one hour before treatment and for 25h. Tubulin is a loading control.

### Accumulation of HIRA in PML NBs depends on SUMO-SIM interactions

We hypothesized that HIRA’s partitioning in PML NBs could be regulated by SUMO-SIM interactions. We first investigated whether HIRA and PML/SUMO could interact together *in cellulo*. Proximity Labelling Assay (PLA) allows the detection of closely interacting protein partners *in situ* at distances below 40nm (Sahin et al. 2016). Using PLA, we detected interaction foci between PML and SUMO2/3 as expected (Sahin et al. 2016), with the number of interaction foci increasing significantly upon IFNβ treatment (Figures 2A-B), which is known to stimulate PML SUMOylation (Stadler et al. 1995). We then assessed the interactions between HIRA and PML or HIRA and SUMO2/3. We could detect a significant interaction between these proteins in presence of IFNβ (Figures 2A-B), accordingly to the accumulation of HIRA in PML NBs. Positive PLA signal between HIRA and SUMO2/3 could either mean that HIRA is SUMOylated or that HIRA interacts with other SUMOylated proteins. The molecular mass of HIRA remained unchanged upon IFNβ treatment of primary cells (Sup. Figure 1D), as previously described (McFarlane et al. 2019), which is not in favor of post-translational modification of HIRA with SUMO groups. In addition, ectopic HIRA mutated on K809, identified as a possible SUMOylated lysine in a SUMO screen (Schimmel et al. 2014; Hendriks et al. 2014), was still recruited in PML NBs similar to the wild-type protein (Sup. Figure 1E), suggesting that at least K809 SUMOylation is dispensable for HIRA recruitment in PML NBs. We thus conclude that HIRA can interact with SUMOylated proteins *in situ*.

**Figure 2.**
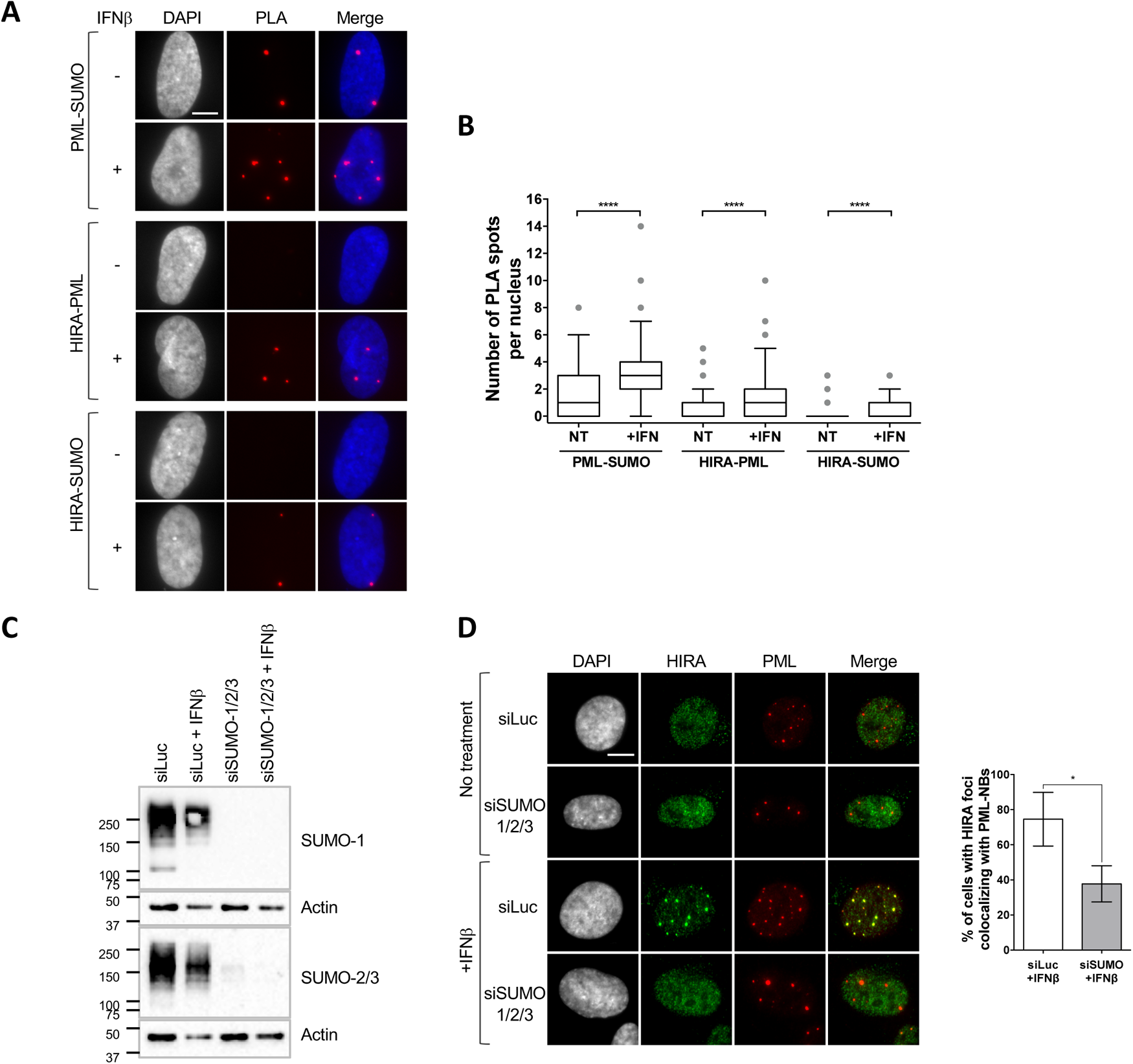
HIRA recruitment to PML NBs is dependent on SUMO proteins. **A.** Fluorescence microscopy visualization of Proximity Ligation Assays (PLA) signals (red) obtained after incubation of anti-PML+anti-SUMO, anti-HIRA+anti-PML or anti-HIRA+anti-SUMO antibodies on BJ cells treated or not with IFNβ at 1000U/mL for 24h. Cell nuclei are visualized by DAPI staining (grey or blue on the merge). Scale bar represents 10μm. **B.** Box-and-whisker plot shows the number of PLA spots detected in cells described in A. In average, 200 nuclei/condition were analyzed from 3 independent experiments. The line inside the box represents the median of all observations. p-values (Mann-Whitney u-test): ****<0,0001. **C.** Western-blot visualization of SUMO-1 and SUMO-2/3 from total cellular extracts of BJ cells treated with 60nM of siRNAs against luciferase or SUMO-1+SUMO-2/3 for 48h and with IFNβ at 1000U/mL during the last 24h. Actin is a loading control. **D.** (left) Fluorescence microscopy visualization of HIRA (green) and PML (red) in BJ cells treated with siRNAs as described in C. Cell nuclei are visualized by DAPI staining (grey). Scale bar represents 10μm. (right) Histograms show quantitative analysis of cells with HIRA localization at PML NBs. Numbers represent the mean of 3 independent experiments (±SD). p-values (Student t-test): *<0,05.

To confirm that SUMOylation of cellular proteins, including PML, is required for HIRA partitioning in PML NBs, we depleted the pool of SUMO1/2/3 by siRNA treatment (Figure 2C). Depletion of SUMOs led to a significant decrease of HIRA accumulation in PML NBs upon IFNβ treatment (Figure 2D). Of note, in absence of SUMOs, PML NBs appear as large aggregates devoid of DAXX (Sup. Figure 2A), reminiscent of the alternative PML NBs structures observed during mitosis, in human embryonic stem cells or in sensory neurons (Corpet et al. 2020). Thus, presence of SUMO proteins, that can undergo LLPS *in vitro* (Banani et al. 2016), seems key to promote partitioning of HIRA. Increasing the pool of free SUMOs by ectopic expression did not trigger HIRA accumulation in PML NBs (Sup. Figure 2B) suggesting that SUMOs need to be conjugated to specific proteins to trigger HIRA partitioning in PML NBs.

To further substantiate the requirements for non-covalent SUMO-SIM interactions in mediating HIRA accumulation in PML NBs, we used the Affimer technology, previously known as Adhiron. Affimers are artificial protein aptamers consisting of a scaffold with two variable peptide presentation loops that can specifically bind with high affinity and high specificity to their binding partners. A recent screen identified several Affimers that inhibit SUMO-dependent protein-protein interactions mediated by SIM motifs (Hughes et al. 2017). We selected the S1S2D5 Affimer that specifically targets both SUMO1 and SUMO2/3-mediated interactions and which possesses a consensus SIM motif (Hughes et al. 2017). Inducibly-expressed S1S2D5-His Affimer showed a nuclear staining with accumulation of the Affimer in PML NBs (Figures 3A-B), as expected for a synthetic peptide that exhibits a SIM domain (Hughes et al. 2017; Banani et al. 2016). S1S2D5-His Affimer expression prevented the accumulation of HIRA in PML NBs upon IFNβ treatment (Figure 3C), without affecting HIRA nor PML protein levels (Sup. Figure 2C). Collectively, our results demonstrate that SUMO-SIM interactions play an important role in the targeting of HIRA in PML NBs in response to IFNβ.

**Figure 3.**
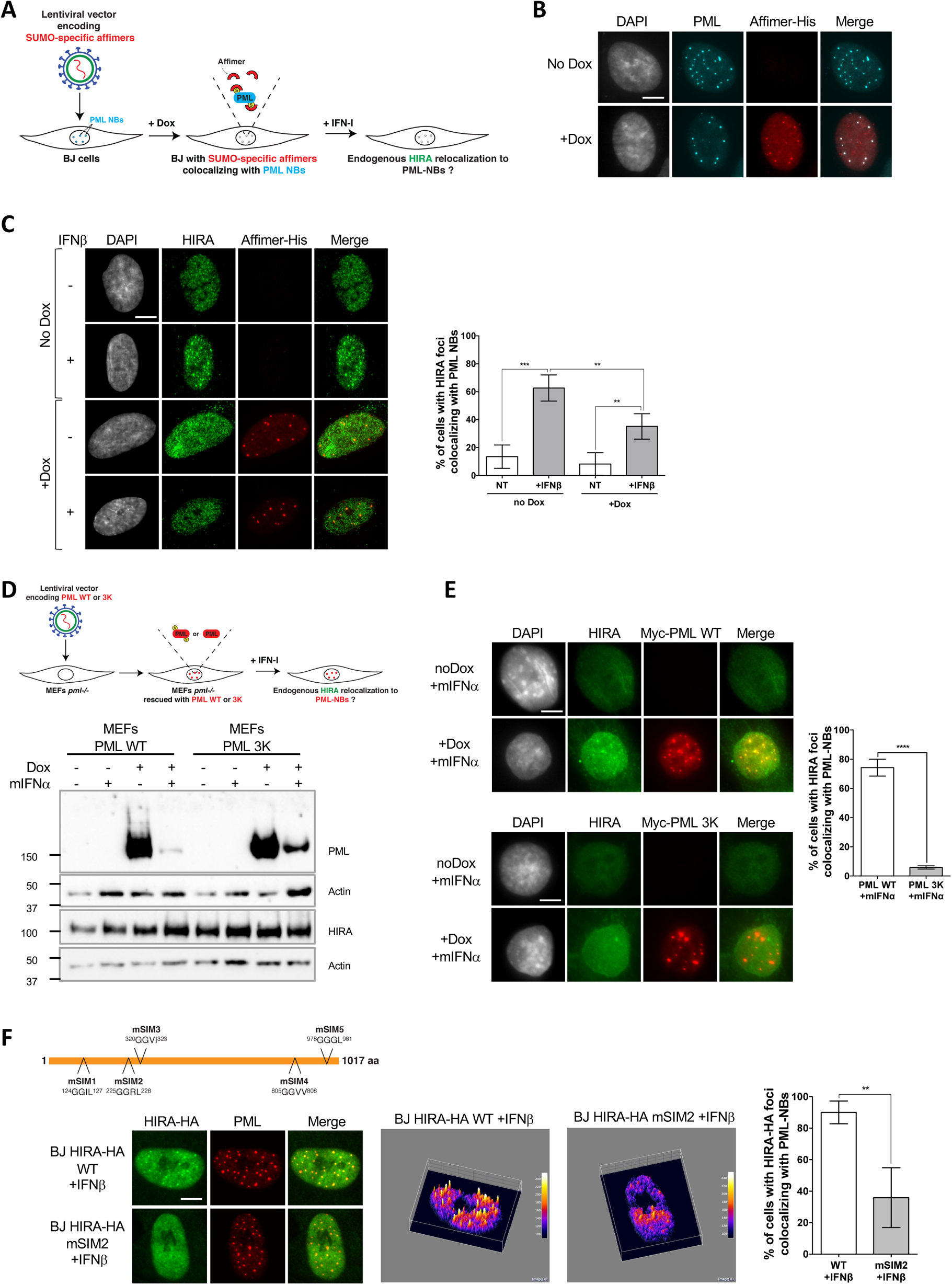
HIRA recruitment to PML NBs relies on SIM-SUMO interactions. **A.** Experimental design to assess SUMO-specific Affimers impact on HIRA relocalization to PML NBs. BJ cells were transduced with a Dox-inducible lentiviral vector encoding for a 6xHis-tagged SUMO-specific S1S2D5 Affimer. When expressed, S1S2D5-His Affimers localize at PML NBs through their interactions with SUMOylated PML. **B.** Fluorescence microscopy visualization PML (cyan) and S1S2D5-His Affimer (red) in transduced BJ cells induced or not with doxycycline at 100nM for 30h. Colocalization of the S1S2D5-His Affimer (red) and PML NBs (cyan) produces white spots. Cell nuclei are visualized by DAPI staining (grey). Scale bar represents 10μm. **C.** (left) Fluorescence microscopy visualization HIRA (green) and S1S2D5-His Affimer (red) in transduced BJ cells induced or not with doxycycline for 30h and treated with IFNβ at 1000U/mL for the last 24h. Cell nuclei are visualized by DAPI staining (grey). Scale bar represents 10μm. (right) Histogram shows quantitative analysis of cells with HIRA localization at PML NBs. Numbers represent the mean of 4 independent experiments (±SD). p-values (Student t-test): **<0,01; ***<0,001. **D.** (top) Experimental design to assess SUMOylated PML requirement for HIRA accumulation to PML NBs. MEFs *Pml*^−/−^ cells were transduced with Dox-inducible lentiviral vectors encoding for Myc-tagged WT or 3K non-SUMOylable PML proteins. Cells were then treated with murine type I IFNα and HIRA localization was observed by fluorescence microscopy. (bottom) Myc-PML proteins expression was verified by western blot analysis of PML from total cellular extracts of MEFs cells describe above. HIRA proteins level was also verified. Actin is a loading control. **E.** (left) Fluorescence microscopy visualization of HIRA (green) and Myc-PML (red) on MEFs *Pml*^−/−^ cells rescued with Myc-tagged WT (top) or 3K (bottom) PML proteins through doxycycline treatment for 24h. Cells were at the same time treated with murine IFNα at 1000U/mL. Cell nuclei are visualized by DAPI staining (grey). Scale bar represents 10μm. (right) Histogram shows quantitative analysis of cells with HIRA localization at ectopic WT or 3K PML NBs. Numbers represent the mean of 3 independent experiments (±SD). p-value (Student t-test): ****<0,0001. **F.** (top) Schematic representation of the localization of the mutations on putative SIM motifs on HIRA protein. (bottom left) Fluorescence microscopy visualization of HIRA-HA (green) and PML (red) in BJ cells stably transduced with HIRA-HA WT or HIRA-HA mSIM2 mutant and treated with IFNβ at 1000U/mL for 24h. Scale bar represents 10μm. Graphics show HA signal intensity of each pixel delimitated within the nuclei in a 3D-surface plot. Higher expression signal appears in yellow to white colors. (bottom right) Histogram shows quantitative analysis of cells with HIRA-HA localization in PML NBs. Numbers represent the mean of 3 independent experiments (±SD). p-value (Student t-test): **<0,01.

PML is known to be mainly SUMOylated on lysines K65, K160 and K490 (Kamitani et al. 1998). Immortalized *Pml−/−* mouse embryonic fibroblasts (MEFs) reconstituted with a doxycyclin-inducible wild-type Myc-tagged version of human PML (Myc-PML WT) or a PML mutated on its three main SUMOylation sites (Myc-PML 3K) were used to investigate the specific requirements for PML SUMOylation in HIRA partitioning (Figure 3D). We first verified that HIRA accumulation in PML NBs was conserved in wild-type but not *Pml−/−* MEFs upon activation of the IFN-I pathway (Sup. Figure 2D). Upon doxycyclin induction, Myc-PML WT or its mutated form were expressed at high levels in *Pml−/−* MEFs (Figure 3D). Despite a diminution in the amount of the ectopic PML proteins following addition of mouse IFNα (Figure 3D), the wild type PML rescued HIRA accumulation in ectopically formed PML NBs unlike the PML 3K (Figure 3E). Of note, super resolution microscopy analyses of PML 3K-expressing MEFs reveal that PML 3K form spherical structures exactly like WT PML (Sahin et al. 2014). These data demonstrate that PML SUMOylation on K65, K160 and K490 is required for HIRA recruitment in PML NBs.

Multivalent interactions between client SIM motifs and SUMOylated lysines on the PML protein are implicated in client recruitment in PML NBs, as shown for DAXX (Banani et al. 2016; Sahin et al. 2014). Using JASSA (Beauclair et al. 2015) and GPS-SUMO (Zhao et al. 2014), we selected a set of 5 putative SIM motifs in HIRA protein sequence and tested whether they were involved in HIRA recruitment in PML NBs by mutating them individually (Figure 3F). Cells expressing the wild-type (WT) tagged version of HIRA (HIRA-HA WT) displayed ectopic HIRA accumulation in PML NBs upon IFNβ treatment (Figure 3F). HIRA-HA mSIM1 and mSIM3 mutants did not show sufficient expression in individual cells to analyze their localization. HIRA-HA mSIM4 and mSIM5 mutants showed a normal accumulation in PML NBs (Sup. Figure 2E). Interestingly, the HIRA-HA mSIM2 showed a significant decrease in its accumulation in PML NBs upon IFNβ treatment (Figure 3F). This data confirms the importance of the SUMO-SIM interaction pathway in general, and particularly at least, the putative SIM2 motif in HIRA for its recruitment in PML NBs. Unfortunately, the levels of HIRA-HA mSIM2 expression remained very low at the cell population level compared to HIRA-HA WT (Sup. Figure 2F), preventing any biochemical analyses. McFarlane et al. previously published that SP100 was required for HIRA localization in PML NBs (McFarlane et al. 2019), and we confirmed those data (Sup. Figures 3A-C). Overall, our data argue for a multistep molecular mechanism involving PML SUMOylation, a putative SIM motif on HIRA and SP100 in the accumulation of HIRA in PML NBs following activation of the IFN-I signaling pathway.

### PML depletion but not HIRA impairs ISGs expression

IFN-I is responsible for the upregulation of hundreds of ISGs as part of the innate immune response participating in the inhibition of virus replication (Shaw et al. 2017). Since HIRA and PML are involved in transcriptional regulation, we investigated whether the partitioning of HIRA in PML NBs upon IFN-I stimulation could play a role in the transcription of ISGs.

We first analyzed the expression of a selected set of ISGs, *MX1*, *OAS1*, *ISG15* or *ISG54* (*IFIT2*), in cells treated with IFNβ for 6 or 24 h. mRNA levels of these ISGs increased strongly after IFNβ treatment, peaking at 6 h of treatment. Transcripts levels then gradually decreased over time, but remained highly expressed at 24 h post addition of IFNβ (Sup. Figure 4A). We then analyzed ISGs mRNA levels in cells depleted of HIRA or PML (Sup. Figure 4B). Depletion of HIRA had no significant impact on ISGs expression at 6 and 24 h of IFNβ stimulation (Figure 4A), consistent with previous reports (Rai et al. 2017; McFarlane et al. 2019). Compared to ISGs transcription climax at 6h post IFNβ, the peak of accumulation of HIRA in PML NBs at 24h post addition of IFNβ (Sup. Figure 3C) suggests that the latter might not be directly involved in ISGs transcriptional upregulation. In contrast, PML depletion led to a significant reduction in ISGs expression both at 6 and 24 h after IFNβ stimulation (Figure 4A). *MX1*, *OAS1*, *ISG15* or *ISG54* mRNA levels were only 27%, 16%, 28% or 35 % of the levels observed in 6h IFNβ-treated cells, respectively. Thus, the results suggest an essential role of PML in the IFN-I-dependent transcriptional upregulation of ISGs, which is independent of HIRA.

**Figure 4.**
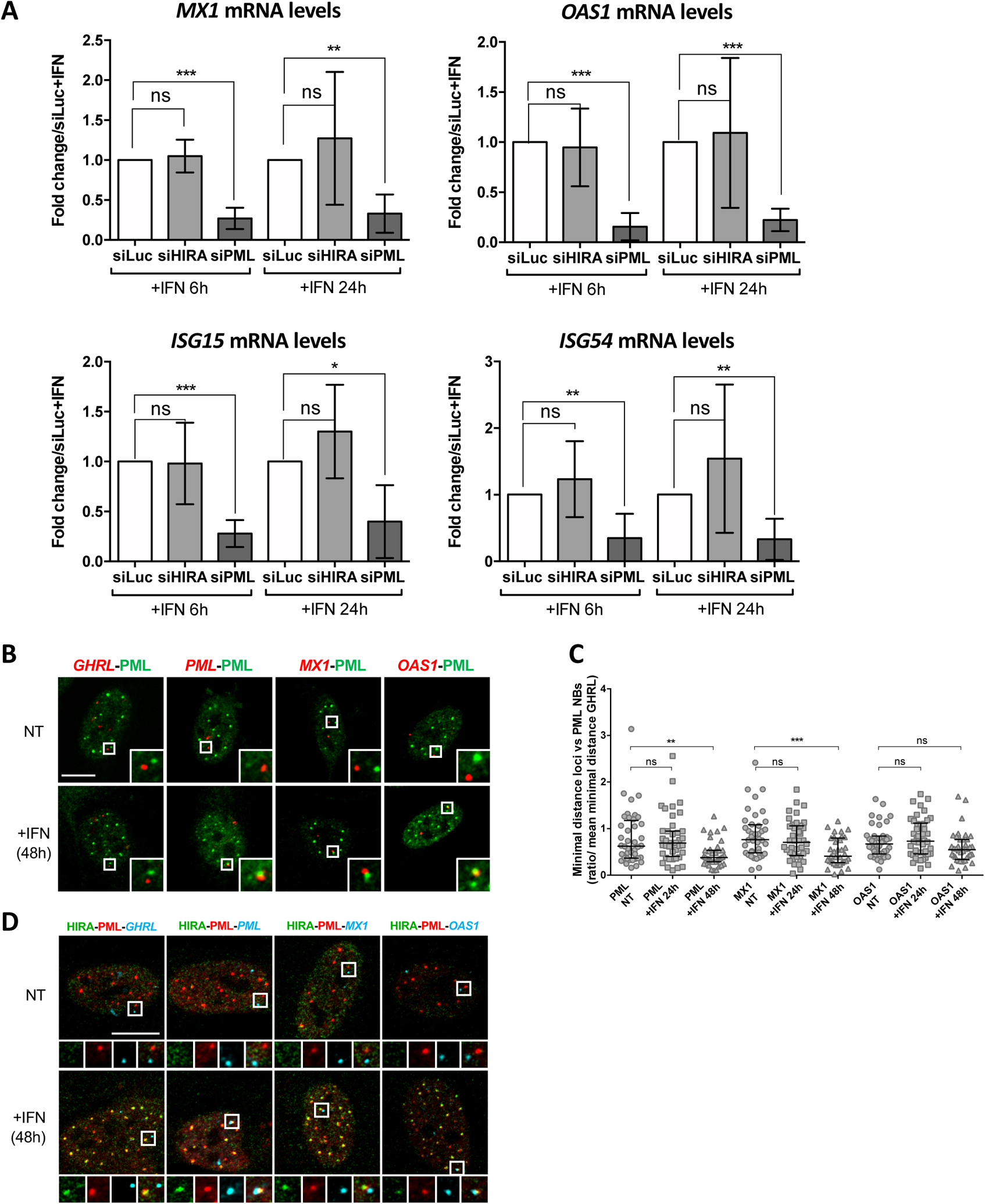
PML is required for ISGs transcription and PML NBs become juxtaposed to ISGs loci. **A.** Histograms show ISGs mRNA relative levels of *MX1*, *OAS1*, *ISG15* and *ISG54* normalized on *GAPDH* mRNA levels of BJ cells treated for 48h with the indicated siRNAs and with IFNβ at 100U/mL for the indicated time. Rationalization was performed on mRNA levels of siLuc+IFN cells. Numbers represent the mean of 3-5 independent experiments (±SD). p-value (Student t-test): *<0,05; **<0,01; ***<0,001; ns: non significant. **B.** Confocal fluorescence microscopy visualization of IF-FISH against PML proteins (green) and *GHRL* control gene locus (red) or *PML*, *MX1* or *OAS1* ISGs loci (red) in BJ cells treated with IFNβ at 1000U/mL for 48h. Insets represent enlarged images (3X) of selected areas and show the relative distance between one PML NB and one gene locus. Scale bar represents 10μm. **C.** Scatter plot shows the ratio of the minimal distance between PML NBs and ISGs loci on the mean minimal distance between PML NBs and *GHRL* control gene locus in nuclei from BJ cells treated or not with IFNβ at 1000U/mL for the indicated time. The line in the middle represents the median of all observations. Results are from one representative experiment out of two experiments and are calculated on an average of 40 nuclei/condition. p-value (Mann-Whitney u-test): **<0,01; ***<0,001; ns: non significant. **D.** Confocal fluorescence microscopy visualization of IF-FISH against HIRA (green) and PML proteins (red) and *GHRL* control gene locus (cyan) or *PML*, *MX1* or *OAS1* ISGs loci (cyan) in BJ cells treated as in B. Insets and scale bar are as in B.

### Interferon-stimulated gene loci are juxtaposed to PML NBs after IFN-I stimulation

PML NBs make direct physical contacts with surrounding chromatin regions and these associations may serve to modulate genome functions and gene expression (Corpet et al. 2020). In the context of the IFNγ inflammatory response, genes within the MHCII locus are located in proximity of PML NBs (Gialitakis et al. 2010). Given the above results, we thought to examine the spatial connection between PML NBs and specific ISG loci. We performed immunostaining of the PML protein, together with fluorescence *in situ* hybridization (immuno-FISH) to detect the *PML*, *MX1*, *OAS1* gene loci in cells treated or not with IFNβ. PML was used as a positive ISG control since previous immuno-trap analyses found a specific interaction between PML NBs and the *PML* gene locus upon IFNα treatment (Ching et al. 2013). To evaluate the specificity of potential spatial changes, we also scored localization of the Grehlin and Obestatin Prepropeptide (*GHRL)* locus, which is not an ISG (Eggenberger et al. 2019) and is localized in heterochromatin regions (Becker et al. 2017). Visual inspection showed an overall closer locus-to-PML NB proximity of *PML*, *MX1* and *OAS1*, but not *GHRL*, in IFNβ-treated cells relative to untreated cells (Figure 4B). To quantify the association of PML NB with ISG loci, we calculated the mean minimal distance (mmd) between each locus and the center of the closest PML NB per nucleus in untreated and treated cells. A decreased distance could be a consequence of the increased number and size of PML NBs upon IFNβ treatment (Sup. Figure 4D). We thus normalized the mmd for the ISGs to the one calculated for the *GHRL* locus. A marked decrease in the mmd of PML NBs with the three loci was scored at 48h, which was significant for *PML* and *MXI* loci, reaching a calculated mmd of 0.67µm and 0.72µm, respectively, as compared to 1.06µm for the *GHRL* locus (Figure 4C). We also confirm the presence of HIRA in PML NBs juxtaposed to ISGs by triple labelling (Figure 4D). The requirement of PML for the acute peak of transcription of ISGs at 6h of IFNβ (Figures 4A), in comparison to the occurrence of the juxtaposition of PML NBs with the ISGs loci at 48h post addition of IFNβ suggests that existing PML NBs are not directly involved in the transcriptional control of ISGs, but rather nucleate at ISGs loci from the PML proteins initially involved in ISGs transcription.

### IFN-I stimulation triggers accumulation of endogenous H3.3 in the 3’ end region of transcribed ISGs

Using a tagged version of H3.3 in MEF cells, previous studies showed an increased and prolonged deposition of ectopic H3.3 in the transcription end sites (TES) region of ISGs upon IFN-I stimulation (Tamura et al. 2009; Sarai et al. 2013). We thus wondered whether PML and HIRA could functionally impact endogenous H3.3 deposition on ISGs using an H3.3-specific antibody previously validated in ChIP (Lee et al. 2019). The amount of H3.3 remained unaffected by 24 h of IFNβ stimulation excluding a putative ISG-like behavior (Sup. Figure 5A). We first investigated H3.3 incorporation on *MX1*, *OAS1* and *ISG54* (*IFIT2*). Three distinct regions of the selected ISGs were analyzed: the promoter region, located just upstream (−120pb) of the transcriptional start site (TSS), the middle of the coding region (mid), and a distal site in the coding region near the TES (see map in Figure 5A). A slight decrease of H3.3 occupancy at promoter regions was measured (Figure 5A). This reduction following IFN stimulation likely reflects transcription-induced nucleosome depletion known to happen for many genes upon stimulation (Workman 2006). Remarkably, IFNβ stimulation induced H3.3 incorporation most noticeably over the distal sites of the coding regions (Figure 5A). This was concomitant with an increase in H3K36me3, a histone mark added by the methyltransferase SETD2, which moves with RNA pol II during transcription (Sup. Figure 5B). Use of a control IgG antibody did not lead to any significant amount of immunoprecipitated DNA (% input) in any of the conditions highlighting the specificity of our ChIP experiment (Sup. Figure 5C). In addition, no change in H3.3 occupancy was observed at an enhancer region known to be enriched with H3.3 (Pchelintsev et al. 2013), underscoring the specificity of H3.3 accumulation in ISGs (Figure 5A). Normalization of H3.3 signal over the total H3 histones signal, which showed no major changes in histone density, confirmed the increased amount of H3.3 at ISGs with a preference for the TES regions (Sup. Figure 5D). This fits with the known replication-independent replacement of canonical H3 histones with H3.3 during transcription (Ahmad and Henikoff 2002; Mito et al. 2005; Workman 2006). No noticeable H3.3 increase was observed at representative mid or TES regions at 6 or 12 h of IFNβ treatment (Sup. Figure 5E). Therefore, H3.3 increased deposition most likely takes place after the peak of ISGs transcription at 6 h of IFNβ (Sup. Figure 4A). Importantly, H3.3 deposition continued to increase for an extended period of time and was even higher at 48 h of IFNβ, suggesting that it could leave a long-lasting chromatin mark on ISGs (Sup. Figure 5E).

**Figure 5.**
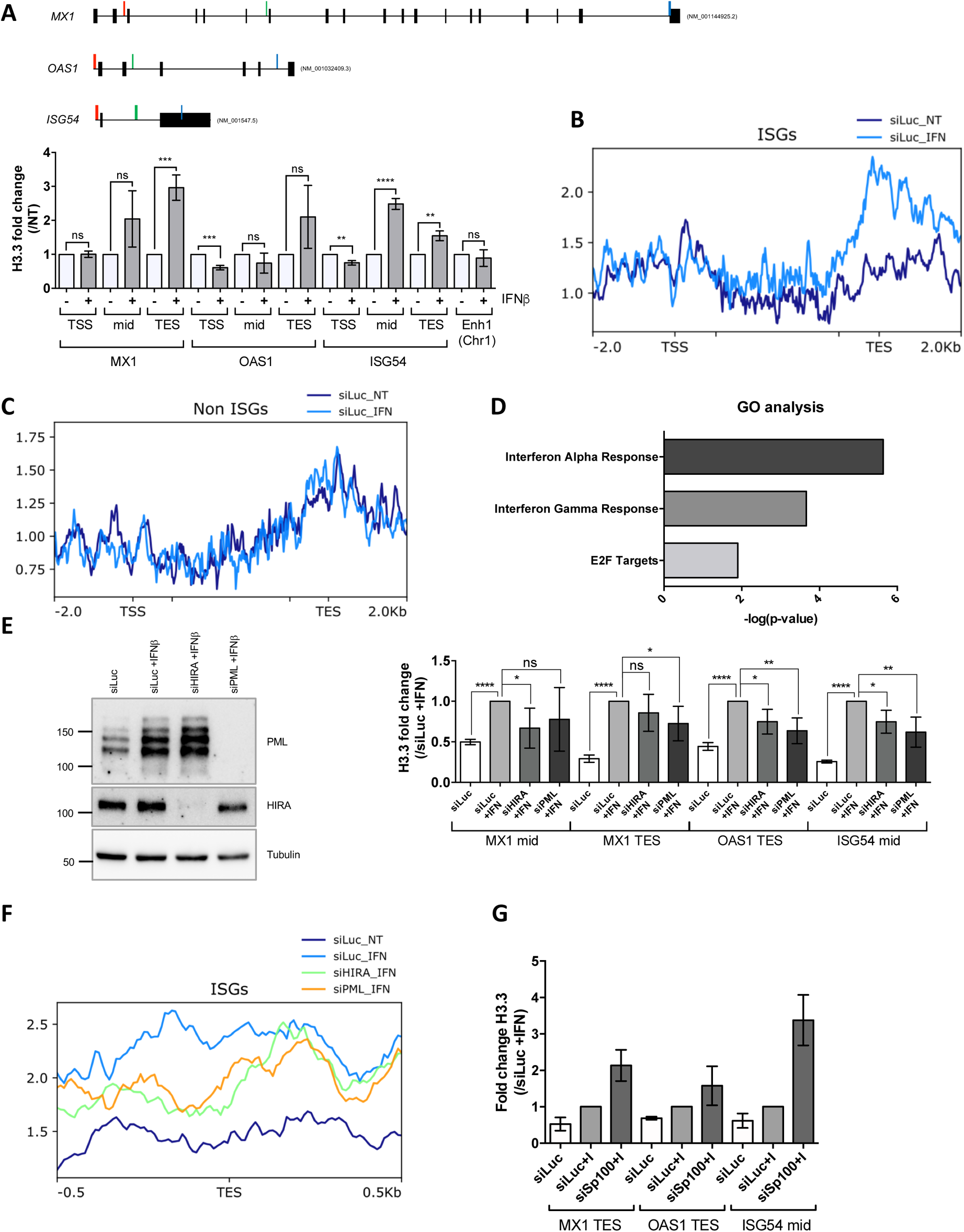
HIRA and PML depletions impair H3.3 enrichment at distal regions of ISGs. **A.** (top) Schematic representation of *MX1*, *OAS1* and *ISG54* gene loci. Localization of primers is marked in color: red, green and blue for primers localized in the Transcription Start Site (TSS), mid or Transcription End Site (TES) region respectively. Black boxes represent exons and lines represent introns. (bottom) Histogram shows H3.3 enrichment fold change obtained through ChIP experiments on BJ cells treated or not with IFNβ at 1000U/mL for 24h. Rationalization was performed on H3.3 enrichment in untreated cells. qPCR was performed on *MX1*, *OAS1* and *ISG54* ISGs TSS, mid and TES regions and on one enhancer region on chromosome 1 (Enh1). Numbers represent the mean of 3 independent experiments (±SD). p-value (Student t-test): **<0,01; ***<0,001; ****<0,0001; ns: non significant. **B.** ChIP-Seq profile of H3.3 enrichment over 49 core ISGs (McFarlane et al. 2019) ranging from −2.0kb to 2.0 kb downstream and upstream of the gene bodies in BJ cells treated as in A. **C.** ChIP-Seq profile of H3.3 enrichment over 49 coding non-ISGs equal in size to core ISGs (McFarlane et al. 2019), ranging from −2.0kb to 2.0 kb downstream and upstream of the gene bodies (regions from TSS to +1000bp and from −1000 to TES being kept unscaled) in BJ cells treated as in A. **D.** Gene Ontology analysis on genes showing the highest differential H3.3 enrichment (log2(Fold Change)>5) in the TES +/- 0.5kb region between IFNβ treated and not treated conditions. **E.** (left) Western blot analysis of HIRA and PML from total cellular extracts of BJ cells treated with the indicated siRNAs for 72h and with IFNβ at 1000U/mL for the last 24h of siRNAs treatment. Tubulin is a loading control. (right) Histogram shows H3.3 enrichment obtained through ChIP experiments on BJ cells treated as on the left panel. Rationalization was performed on H3.3 enrichment in siLuc +IFN treated cells. qPCR was performed on *MX1* mid and TES regions, *OAS1* TES region and *ISG54* mid region. Numbers represent the mean of 4 independent experiments (±SD). p-values (Student t-test): *<0,05; **<0,01; ****<0,0001; ns: non significant. **F.** ChIP-Seq profile of H3.3 enrichment over 49 core ISGs (McFarlane et al. 2019) ranging from −0.5kb to 0.5kb downstream and upstream of the TES in BJ cells treated as in E. **G.** Histogram shows H3.3 enrichment obtained through ChIP experiments on BJ cells treated with the indicated siRNAs for 72h and with IFNβ at 1000U/mL for the last 24h of siRNAs treatment. Rationalization was performed on H3.3 enrichment in siLuc +IFN treated cells. qPCR was performed on *MX1* and *OAS1* TES regions and on *ISG54* mid region. Numbers represent the mean of 3 technical replicates out of 2 independent experiments (±SD).

We then performed ChIP-Seq analysis for endogenous H3.3 on cells treated or not with IFNβ for 24 h. We first examined H3.3 enrichment over the gene bodies of a published panel of equivalent sized ISGs or non-ISGs (McFarlane et al. 2019). The levels of H3.3 significantly increased on ISGs in IFNβ-treated cells with a clear bias towards the TES regions of the genes (Figure 5B). In contrast, no significant difference of H3.3 enrichment could be observed across the non-ISGs (Figure 5C). We selected genes with the highest difference in H3.3 enrichment at the TES region between IFNβ treated and non-treated cells, and performed Gene Ontology (GO) analysis. GO analysis showed a clear enrichment in genes involved in IFNα and IFNγ response comforting the specific enrichment of H3.3 on the TES region of ISGs as a prolonged response to an IFN-I stimulus (Figure 5D). To evaluate the identity of H3.3-enriched genes in an unbiased manner, we performed an independent GO analysis on all genes found in the ChIP peak calls. This yielded similar results with IFNα and IFNγ responses being the most highly significant GO terms (Sup. Figure 5F). Thus, these findings establish that IFN-I triggers a specific long-lasting H3.3 deposition on ISGs following IFN-I stimulus.

### H3.3 deposition on ISGs is impaired upon HIRA or PML depletion but is independent of the localization of HIRA in PML NBs

We next wondered whether HIRA and/or PML was essential for H3.3 deposition at ISGs. Cells were depleted of HIRA or PML (Figure 5E, left) and treated with IFNβ for 24 h, before performing ChIP on H3.3. Absence of HIRA or PML led to a modest but consistent decrease in H3.3 at mid or TES regions of selected ISGs, suggesting the implication of these two proteins for the long-lasting H3.3 deposition on ISGs (Figure 5E, right). ChIP-Seq analysis confirmed a modest, but still significant, decrease in the loading of H3.3 at the TES on the panel of ISGs (p-value = 4,76e-03 for HIRA knock-down (KD) or 1.262e-03 for PML KD, as assessed by a paired Student’s t-test) (Figure 5F, Sup. Figure 5G). Representative *STAT1* and *GCH1* genes, confirmed the deficit in H3.3 loading at the TES region of ISGs in the absence of HIRA or PML (Sup. Figure 5H). We thus conclude that HIRA and PML both contribute to the increased long-lasting H3.3 deposition at the TES region of ISGs following the transcriptional peak associated to IFN stimulus.

We then investigated if the accumulation of HIRA in PML NBs was a prerequisite for the increased deposition of H3.3 on ISGs. The low level of expression of the HIRA-HA mSIM2 mutant, that does not accumulate in PML NBs following IFN-I stimulus (Sup. Figure 2F), unfortunately precluded its use for biochemical analyses. We therefore chose to deplete SP100 that also strongly impairs HIRA recruitment in PML NBs (Sup. Figure 3 and (McFarlane et al. 2019)). SP100 depletion did not prevent H3.3 loading at the ISGs TES upon IFN-β stimulation, but on the contrary increased it (Figure 5G). These data suggest that although both HIRA and PML are essential for the long-lasting deposition of H3.3 on the TES of ISGs following an acute IFN-I stimulus, the accumulation *per se* of HIRA in the PML NBs serves a different purpose.

### PML NBs act as storage sites to buffer HIRA according to histones availability and dynamics

Quantification of nucleoplasmic levels of HIRA outside PML NBs showed a significant decrease after IFNβ treatment (Figure 6A). Similarly, ectopic expression of HIRA led to its accumulation in PML NBs without the need of IFN-I stimulation (Figure 6B), suggesting that PML NBs could help buffering an excess of HIRA proteins. H3.3-H4 deposition/recycling on chromatin is tightly regulated by the HIRA complex (Martire and Banaszynski 2020). We thus reasoned that an increase of the nucleoplasmic pool of H3.3-H4 could modulate HIRA accumulation in PML NBs upon IFN-I treatment. We generated human primary BJ cells expressing an inducible HA-tagged form of H3.3 (BJ eH3.3i). Treatment with doxycyclin triggered a strong, yet highly variable, expression of eH3.3 (Figure 6C, Sup. Figures 6A-B), consistent with the polyclonal nature of the BJ eH3.3i. We verified that doxycyclin did not impact HIRA localization in absence of IFNβ and that IFNβ alone triggered a normal accumulation of HIRA in PML NBs in these cells (Sup. Figure 6A). Addition of doxycyclin that triggers H3.3 overexpression did not significantly alter HIRA accumulation in PML NBs upon IFNβ on a population level (Sup. Figure 6A). However, close examination of individual cells showed an impaired accumulation of HIRA in PML NBs in cells with a strong overexpression of eH3.3 (Figure 6C). This was not observed in low eH3.3 expressing cells, in which HIRA was detected in PML NBs together with eH3.3 (Figure 6C). Quantification of the mean eH3.3 nuclear fluorescence intensity showed that it was low in average in nuclei showing accumulation of HIRA in PML NBs upon IFNβ, while a significant shift to higher eH3.3 nuclear intensities was observed in nuclei showing absence of HIRA in PML NBs, underscoring a strong antagonism between HIRA detection in PML NBs and high expression of eH3.3 (Figure 6D). Similar results were obtained in human primary lung fibroblasts (Sup. Figure 6C). Therefore, presence of an excessive H3.3 pool in the nucleoplasm prevents HIRA accumulation in PML NBs upon IFN-I stimulation.

**Figure 6.**
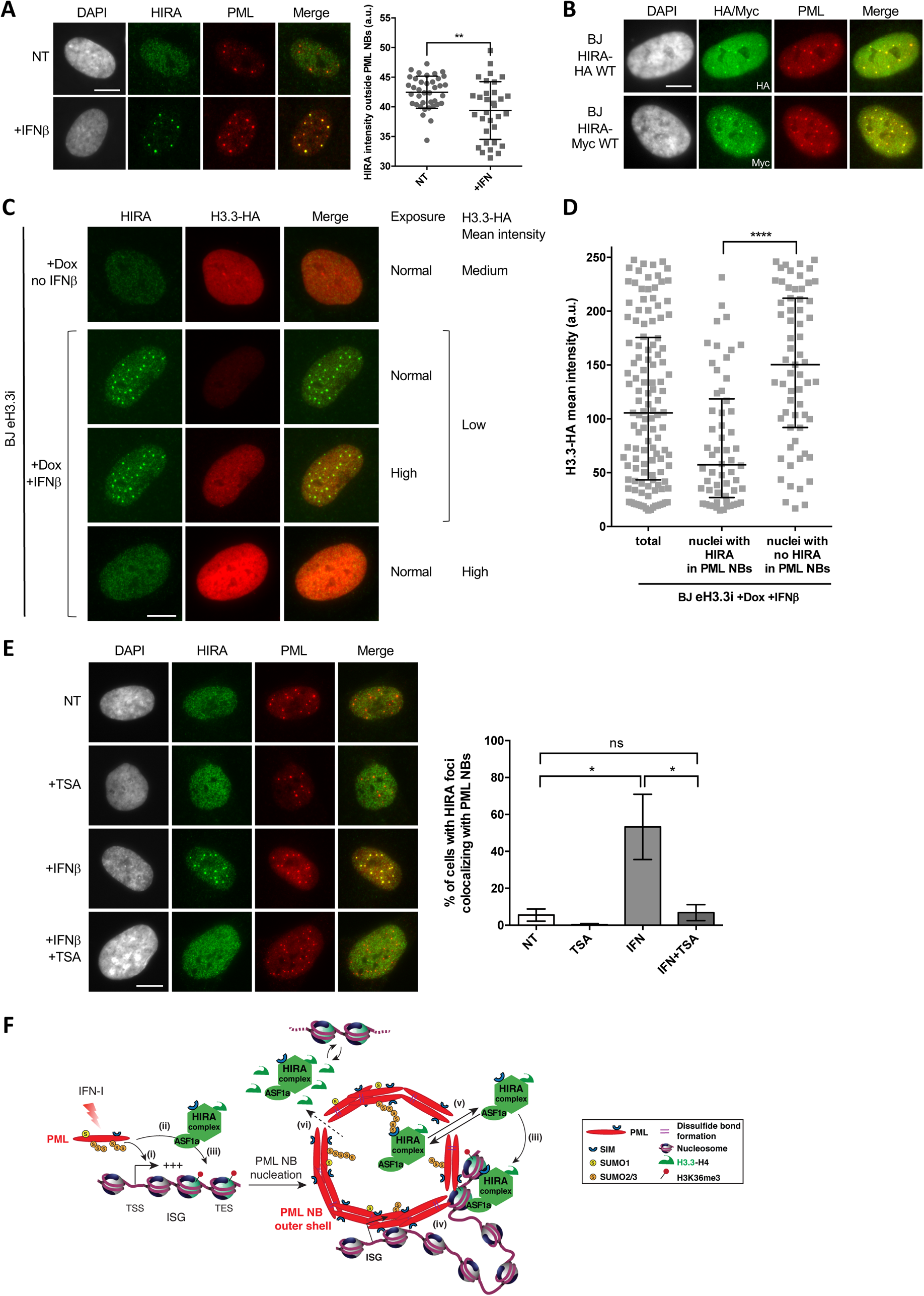
HIRA accumulation in PML NBs depends on the availability of a nucleoplasmic H3.3-H4 pool and on chromatin compaction. **A.** (left) Fluorescence microscopy visualization of HIRA (green) and PML (red) in BJ cells treated with IFNβ at 1000U/mL for 24h (+IFNβ) or left untreated (NT). Cell nuclei are visualized by DAPI staining (grey). Scale bar represents 10μm. (right) Histogram shows quantification of HIRA mean nuclear fluorescence intensity outside PML NBs (a.u : arbitrary units) in nuclei from 5 independent experiments. p-values (Mann-Whitney u-test): **<0,01 **B.** Fluorescence microscopy visualization of HA or Myc (green) and PML (red) in BJ HIRA-HA WT and BJ HIRA-Myc WT cells. Cell nuclei are visualized by DAPI staining (grey). Scale bar represents 10μm. **C.** Fluorescence microscopy visualization of HIRA (green) and H3.3-HA (red) in BJ eH3.3i cells treated with doxycyclin and with or without IFNβ at 1000U/mL for 24h. High exposure indicates a lane where H3.3-HA signal was specifically increased in order to show H3.3-HA localization in PML NBs without saturating the signal in cells with higher expression. Scale bar represents 10μm. **D.** Quantification of nuclear H3.3-HA intensity levels in BJ eH3.3i cells treated as in C. Mean H3.3-HA intensity levels were calculated on a pool of n=121 nuclei from 3 independent experiments. Nuclei were then separated on basis of accumulation of HIRA in PML NBs (nuclei with HIRA in PML NBs, n=58) or without it (nuclei with no HIRA in PML NBs, n=63) and mean H3.3-HA intensity was plotted for each category. Bars represent median with interquartile range. p-values (Mann-Whitney u-test): ****<0,0001. **E.** (left) Fluorescence microscopy visualization of HIRA (green) and PML (red) in BJ cells treated with IFNβ at 1000U/mL for 24h (+IFNβ) or left untreated (NT). TSA at 2μM was added or not 1h before IFNβ. Cell nuclei are visualized by DAPI staining (grey). Scale bar represents 10μm. (right) Histogram shows quantitative analysis of cells with HIRA localization at PML NBs. Numbers represent the mean of 3 independent experiments (±SD). p-values (Student t-test): *<0,05; ns: non significant. **F.** Model for the dual role of PML/PML NBs in inflammatory response. At early time points after IFN-I treatment, (i) PML is required for ISGs transcription and (ii) it could help to load HIRA on ISGs (McFarlane et al. 2019), participating in H3.3 deposition. (iii) While HIRA depletion does not affect ISGs transcription *per se*, it could participate in H3.3 deposition/recycling at ISGs, a function which does not seem to require its accumulation in PML NBs. (iv) PML neonucleation would mediate juxtaposition of PML NBs with ISGs at late times after IFN-I treatment which could help to keep a memory of the physiological state of the cell. (v) In addition, PML NBs play a second independent role by buffering the pool of HIRA complex available in the nucleus. (vi) Increase in the soluble pool of H3.3-H4 dimers or increase in histone exchange/dynamics upon chromatin decompaction could force HIRA out of PML NBs.

Finally, we wondered if increasing histone dynamics could also prevent HIRA accumulation in PML NBs by forcing its requirement on chromatin. We therefore treated BJ cells with trichostatin A (TSA), a drug that selectively inhibits class I and II mammalian histone deacetylases, inducing chromatin decompaction and therefore increasing histone exchange on chromatin (Nozaki et al. 2017). Interestingly, TSA treatment significantly impaired the accumulation of HIRA in PML NBs seen under IFNβ stimulation (Figure 6E), without noticeable change in HIRA levels (Sup. Figure 6D). Thus, TSA-induced hyperacetylation of chromatin seems to redirect HIRA from PML-NBs, possibly to ensure H3.3 deposition/recycling on the hyperactive decondensed chromatin.

## Discussion

There have been considerable efforts in defining the multiple roles of PML and PML NBs in the recent years including in chromatin dynamics. After having dissected the molecular mechanisms responsible for HIRA accumulation in PML NBs upon IFN-I treatment, we investigated the functional role of the PML NBs-HIRA-H3.3 axis in inflammatory response. Our work underscores two independent roles for PML/PML NBs in (1) regulating the transcriptional status of ISGs and the incorporation of H3.3 at these loci and in (2) acting as storage centers to modulate HIRA complex nucleoplasmic availability upon inflammatory stress.

### HIRA accumulation in PML NBs upon inflammatory stresses is dependent on functional SUMO-SIM interactions

While senescence was the first stress shown to induce accumulation of HIRA complex in PML NBs (Zhang et al. 2005; Banumathy et al. 2009; Rai et al. 2011; Jiang et al. 2011), IFN-I signaling pathway was recently shown to be responsible for similar behavior of HIRA upon a viral infection (Rai et al. 2017; Cohen et al. 2018; McFarlane et al. 2019). Here, we first extend and corroborate these findings by showing that various inflammatory stresses, including TNFα, or a synthetic dsRNA (PolyI:C), can also mediate HIRA recruitment in PML NBs. We show that HIRA partitioning in PML NBs is mediated by SUMO-SIM interactions, that can be inhibited with specific Affimers. Our data using *Pml^−/−^* MEFs reconstituted with PML WT or PML 3K underline the importance of PML main SUMOylation sites in recruiting HIRA complex. Remarkably, SIM-containing clients, as exemplified with DAXX (Sahin et al. 2014), are favored for partitioning in endogenous PML NBs containing a higher valency of SUMO sites compared to SIM sites (Banani et al. 2016). Here, we identified a putative SIM motif on HIRA sequence that participate in its recruitment in PML NBs. Interestingly, the VLRL SIM motif identified is followed by a Serine in the position 229 (S229). Phosphorylation adjacent to SIM motifs can lead to an increased affinity towards SUMO1 lysine residues (Cappadocia et al. 2015). Other post-translational modifications such as phosphorylation could thus be important in regulating HIRA partitioning by changing the affinity between HIRA and SUMOylated PML proteins/partners. Of note, glycogen synthase kinase 3 β (GSK-3β) mediated-phosphorylation of HIRA on S697 was suggested to drive HIRA accumulation in PML NBs upon senescence entry (Ye et al. 2007). Thus, HIRA accumulation in PML NBs after IFN-I treatment mechanistically relies on SUMO-SIM interactions. Whether this is mediated by direct/indirect interactions with SUMOylated PML or SP100 ((McFarlane et al. 2019) and this study), remains to be investigated. Nevertheless, the use of SUMO-specific Affimers opens interesting avenues to interfere with client recruitments in PML NBs including HIRA.

### PML regulates ISGs transcription and PML NBs associate with ISGs loci

We next investigated the functional role of the accumulation of HIRA in PML NBs by first focusing on the impact of HIRA and PML depletion in ISGs transcription and H3.3 deposition. While HIRA binding to ISGs is increased after IFN-I (McFarlane et al. 2019; Rai et al. 2017), its depletion did not affect ISGs expression, consistent with previous reports (McFarlane et al. 2019; Rai et al. 2017). In contrast, our data underscore the importance of the PML protein for the initial burst of transcription of ISGs at 6 h of IFNβ stimulation. This is consistent with both the association of PML NBs with transcriptional sites after IFNβ stimulation, as shown by visualization of nascent transcripts (Fuchsová et al., 2002) and consistent with the role of PML in ISGs induction following viral infection (Alandijany et al., 2018). PML proteins could be recruited to transcriptionally active ISGs by a specific, yet to be defined, protein-protein interaction. Previous studies showed that the nuclear DNA helicase II (NDH II), which is essential for gene activation, relocalizes in PML NBs in a transcription-dependent manner (Fuchsová et al. 2002). The authors suggested PML NBs could play a role in the transcriptional regulation of ISGs attached to PML NBs, although this was not investigated (Fuchsová et al. 2002). Here, by using immuno-FISH, we demonstrate for the first time a juxtaposition of a subset of ISG loci with PML NBs at late time-points of IFNβ stimulation. This data underscores the likely importance of a PML NBs-gene locus association as a marker of the physiological state of the cell. Since, PML targeting at specific gene loci is sufficient to induce de novo formation of PML NBs (Brouwer et al., 2009; Chung et al., 2011; Erdel et al., 2020; Kaiser et al., 2008; Wang et al., 2018), we hypothesize that chromatin-bound PML proteins involved in ISGs transcription could then act as seeds to mediate neonucleation of new PML NBs at ISG loci (see model in Figure 6F). Alternatively, movement of ISGs to preexisting PML NBs could still be at play to explain this closer association upon IFNβ stimulation. The use of ALaP-Seq method to map chromatin regions proximal to PML NBs (Kurihara et al., 2020) could nicely complement the immuno-FISH approach to identify genome-wide changes in PML-NBs-chromatin associations upon IFN-I stimulation.

### H3.3-induced deposition in the TES region of transcribed ISGs is partially mediated by HIRA and PML

We then investigated the role of PML and HIRA in the H3.3-mediated deposition on ISGs. Endogenous H3.3 deposition shows a strong preference for the ISGs TES regions, consistent with previous reports obtained in mouse cells overexpressing exogenous H3.3 (Tamura et al. 2009; Sarai et al. 2013). Also, our data highlight a long-lasting deposition of H3.3 up to 48h after IFN-I stimulation, well beyond the peak of transcription of the ISGs. Deposition of endogenous H3.3 was reduced in the absence of PML consistent with the role of PML NBs in targeting H3.3 to chromatin (Delbarre et al. 2013) and in line with the role of PML in chromatinization of latent viral genomes (Cohen et al. 2018). Since PML depletion impairs transcription of ISGs (Figure 4A) and (Alandijany et al. 2018), it could as well indirectly affect H3.3 deposition at TES regions, which has been shown to be linked to the transcriptional activity of ISGs *per se* (Sarai et al. 2013). PML has been shown to be implicated in the loading of HIRA on ISGs (McFarlane et al. 2019), anticipating a role of a PML-HIRA axis in H3.3 deposition on these loci. HIRA depletion indeed mildly impaired H3.3 deposition at ISGs TES regions, which is consistent with its known function in H3.3-nucleosome assembly. HIRA interacts with RNA pol II and also with H3K36me3 (Torné et al. 2020; Ray-Gallet et al. 2011), a histone mark added by the methyltransferase SETD2, which moves with RNA pol II during transcription. In MEFs, HIRA has been shown to interact with WHSC1, another H3K36me3 methyltransferase that recruits HIRA for prolonged H3.3 deposition on ISGs (Sarai et al. 2013). Since H3K36me3 was also found to be enriched at the TES part of ISGs in IFNβ treated cells (Sup. Figure 5B), one could anticipate a similar recruitment of HIRA via an H3K36methyltransferase/H3K36me3 axis. Of note, HIRA impact on H3.3 levels could also be explained by its role in H3.3 recycling during ISGs transcription (Torné et al. 2020). Since H3.3 deposition is only moderatly affecting following HIRA depletion, it is not unlikely that other H3.3 chaperones could compensate for the absence of HIRA as already observed for viral genomes chromatinization (Cohen et al. 2018). Indeed, the DAXX/ATRX complex, which localizes constitutively in PML NBs, could participate in H3.3 deposition on ISGs, although its activity is mainly associated to chromatin silencing. Alternatively, the remodeling protein CHD2 has been shown to be recruited to the promoters of myogenesis genes to incorporate H3.3 (Harada et al. 2012, Siggens et al., 2015). It could be interesting to see if CHD2 is recruited to PML NBs, and if it participates to H3.3 deposition at ISGs loci after IFN-I induction. Nonetheless, these data indicate the importance of a PML-HIRA axis to regulate H3.3 dynamics on ISGs loci.

The role of the prolonged H3.3 deposition on ISGs can be multiple. First, this long-lasting mark could contribute to the acquisition of a functional IFN response memory. Indeed, H3.3 was shown to mediate memory of an active state upon nuclear transfer in *Xenopus laevis* (Ng and Gurdon 2007). In addition, in MEFs, IFNβ stimulation creates a transcriptional memory of a subset of ISGs, which coincides with acquisition of H3.3 and H3K36me3 on chromatin (Kamada et al. 2018). A second stimulation with IFNβ allows a faster and greater transcription of so called “memory ISGs”, which is dependent on H3.3 deposition during the first stimulation phase (Kamada et al. 2018). In HeLa cells, PML was shown to be required for the stronger re-expression of HLA-DRA after IFNγ restimulation, a locus that remained juxtaposed to PML NBs after transcription shut-off (Gialitakis et al. 2010). Second, H3.3 deposition may also serve to directly regulate ISGs expression. In mouse cells, H3.3 was found to be phosphorylated on Serine 31 on macrophages-induced genes following bacterial lipopolysaccharide stimulation, a post-translational mark serving as an ejection switch for the ZMYND11 transcriptional repressor, and allowing the transcriptional amplification of the target genes (Armache et al. 2020). Whether H3.3S31P is increased on ISGs upon IFN-I remains to be investigated.

Given the accumulation of HIRA in PML NBs upon IFNβ stimulation, an important question was to address whether this accumulation was required for HIRA function in H3.3 deposition/recycling at ISGs. Our ChIP analysis after SP100 depletion, which prevents HIRA localization in PML NBs under IFN-I stimulation is impaired, did not show any decrease of H3.3 enrichment at ISGs loci but rather an increase. This result suggests that HIRA accumulation in PML NBs is not a prerequisite for H3.3 deposition on ISGs, at least following a first wave of IFN-I stimulation, and might rather serve as a nuclear depot for subsequent activities.

### PML NBs as nuclear depot centers to regulate nucleoplasmic and chromatin-bound HIRA levels

Importantly, our study unveils a second aspect of the PML NBs-HIRA interplay, with PML NBs seemingly acting as depot centers to regulate the pool of nucleoplasmic/chromatin-bound HIRA, independently of their roles in ISGs transcription/H3.3 deposition. We first show that HIRA intensity level in the nucleus, outside PML NBs, decreases upon IFN-I treatment, while steady state HIRA protein amount remains unchanged (our study and (Rai et al. 2017)). Acute stress could thus induce changes in HIRA requirements in the nucleus, to retarget it to specific chromatin loci, leaving a pool of “unemployed” HIRA, which then accumulates in PML NBs and could be released later for further tasks. Hence, it is interesting to mention that overexpression of an ectopic HIRA is also sufficient to induce its accumulation in PML NBs in untreated cells ((Ye et al. 2007) and this study), underscoring the fact that an excess of HIRA protein is indeed buffered in PML NBs.

Although SUMO-SIM interactions play an essential role in mediating HIRA accumulation in PML NBs, our study unveils the availability of free H3.3 as a novel important parameter controlling this accumulation. Indeed, overexpression of a pool of free soluble H3.3 in the nucleoplasm impairs HIRA accumulation in PML NBs upon IFN-I treatment, possibly by driving HIRA outside PML NBs to handle this pool of H3.3-H4 histones. Similarly, TSA-induced hyperacetylation, which results in global decompaction of chromatin and increased histone exchange/dynamics (Nozaki et al. 2017), also strongly impairs HIRA localization in PML NBs upon IFN-I treatment.

Accordingly, we propose a role for PML/PML NBs in regulating ISGs transcription and H3.3 deposition as follows (Figure 6F): PML is required for ISGs transcription promoting H3.3 loading on these genes. In addition, PML could serve as a plateform to load HIRA on ISGs (McFarlane et al. 2019). While HIRA depletion does not affect ISGs transcription *per se*, it could participate in H3.3 deposition/recycling at ISGs, a function which does not seem to require its accumulation in PML NBs. In addition, PML NBs plays a role of ‘nuclear storage center’ for HIRA complex, to buffer its availability in the nucleoplasm, and thus possibly regulating its activity (Figure 6F). As mentioned above, H3.3 deposition at ISGs after a first stimulus allows faster and greater transcription of ISGs upon restimulation (Kamada et al. 2018). Juxtaposition of PML NBs with ISGs at late times after IFN-I treatment could help to keep a memory of the physiological state of the cell. PML would remain in close proximity to ISGs to regulate them upon a second wave of IFN-I stimulation and HIRA accumulation in PML NBs could also be a mean for the cell to make the chaperone complex available much faster in case of a second inflammatory wave.

In conclusion, our study highlights two important functional and independent roles for PML NBs in the inflammatory response, which add to their pivotal involvment in various stress responses.

## Methods

### Cell lines and lentiviruses production

Human BJ primary foreskin fibroblasts (ATCC, CRL-2522), human IMR90 fetal lung fibroblasts (ATCC, CCL-186), human HEK 293T embryonic kidney cells (Intercell, AG) and mouse MEFs embryonic fibroblasts *Pml*^−/−^ (from Dr. Lallemand-Breitenbach) were cultivated in DMEM medium (Sigma-Aldrich, D6429) containing 10% of fetal calf serum (FCS) (Sigma-Aldrich, F7524), 1% of penicillin/streptomycin (Sigma-Aldrich, P4458), at 37°C under 5% CO2 and humid atmosphere. Drugs and molecules used for cell treatments are described in **Sup. Table 1** (duration is mentioned in the main text). BJ, MEFs or IMR90 cells stably expressing transgenes were obtained by lentiviral transduction as in (Cohen et al. 2018). Transduced cells were then selected 24h later by adding the appropriate selective drug (puromycin (Invivogen, ant-pr) at 1μg/mL or blasticidin (Invivogen, ant-bl) at 5μg/mL).

### Plasmids

Tat-S1S2D5-Flag-His Affimer (Tat: nuclear localization sequence), obtained by PCR using pcDNA5-Tat-S1S2D5-Flag-His as template (graciously sent by Dr. David J. Hughes (Hughes et al. 2017)), was cloned in puromycin resistant pLVX-TetOne plasmid.

HIRA WT, obtained by RT-PCR from HeLa cells, was cloned in blasticidin resistant pLentiN plasmid with addition of HA or Myc tag in the C-terminus. HIRA-HA mSIM and K809G mutants were obtained by site-directed mutagenesis using QuickChange Lightning Site-directed Mutagenesis kit (Agilent Technologies, #210518). SIM motifs are characterized by a group of hydrophobic amino acids ((V/I/L)x(V/I/L)(V/I/L) or (V/I/L)(V/I/L)x(V/I/L)). HIRA mSIM mutant sequences are the following: mSIM1: aa ^124^VSIL^127^ mutated in ^124^GGIL^127^, mSIM2: aa ^225^VLRL^228^ mutated in ^225^GGRL^228^, mSIM3: aa ^320^LLVI^323^ mutated in ^320^GGVI^323^, mSIM4: aa ^805^VVVV^808^ mutated in ^805^GGVV^808^, mSIM5: aa ^978^VVGL^981^ mutated in ^978^GGGL^981^.

Myc-PML1 WT and 3K mutant, obtained by PCR using pLNGY-PML1 and pLNGY-PML1.KKK as template (kind gift by Dr. Roger Everett), were cloned in puromycin resistant pLVX-TetOne plasmid. H3.3-SNAP-HA3 obtained by PCR using pBABE-H3.3-SNAP-HA3 as template (kind gift by Dr. Lars Jansen) was cloned into puromycin-resistant pLVX-TetOne plasmid with EcoRI restriction enzyme.

### siRNAs

BJ cells were transfected with 60nM of human siRNA for different timings (indicated in the main text for each experiment) using Lipofectamine RNAiMax reagent (Invitrogen, 13778-075) and Opti-MEM medium (Gibco, 31-985-070). siRNAs used and their sequences are summarized in **Sup. Table 2**. siSUMO1 and siSUMO-2/3, were co-transfected into BJ cells at 30nM each.

### Antibodies

All the primary antibodies used in this study, together with the species, the reference and the dilutions for immunofluorescence and western blotting, are summarized in **Sup. Table 3**.

### Immunofluorescence (IF)

Immunofluorescence was performed as in (Corpet et al. 2014) (see **Sup. Table 3** for antibodies dilution). HIghly cross-absorbed goat anti-mouse or anti-rabbit (H+L) Alexa-488, Alexa-555 or Alexa-647 (Invitrogen) were used as secondary antibodies. Cells were then incubated in DAPI (Invitrogen Life Technologies, D1306) diluted in PBS at 0,1μg/mL for 5 minutes at RT°C. Coverslips were mounted in Fluoromount-G (SouthernBiotech, 0100-01) and stored at 4°C before observation.

### Proximity Ligation Assay (PLA)

Proximity Ligation Assays were performed with the Duolink In Situ Red Starter Kit Mouse/Rabbit (Sigma-Aldrich, DUO92101). Cells on coverslips were fixed in 2% PFA for 12 minutes at RT°C and then permeabilized in PBS 0,2 % Triton X-100 for 5 minutes at RT°C. Cells were then treated according to the manufacturer’s instructions (see **Sup. Table 3** for dilutions of primary antibodies). Coverslips were mounted in Duolink In Situ Mounting Medium with DAPI and stored at 4°C before observation.

### Immunofluorescence - Fluorescence in situ hybridization (IF-FISH)

FISH probes were generated from different BACs: RP11-438J1, RP11-185E17, RP11-120C17 and RP11-134B23 BAC clones for GHRL, PML, MX1 and OAS1, respectively. Briefly, 1μg of BAC were incubated for nick-translation with 4.3 ng of DNAse I (Roche, 104159), 7U of DNA polymerase (Promega, M2051), dithiothreitol (DTT) at 10μM, dATP, dTTP and dGTP at 40μM each (Thermo Scientific, R0141/R0161/R0171), dCTP at 10μM (Thermo Scientific, R0151) and Cy3 labelled dCTP at 10μM (Cytiva, PA53021). Nick-translation was performed for 3 h at 15°C and stopped by an incubation at 72°C for 10 minutes. Size of generated probes were verified on agarose gel. Probes were then mixed with 20μg of COT Human DNA (Roche, 11 581 074 001) and 79μg of Salmon sperm DNA (Invitrogen, 15632-011). Volume was completed with TE buffer (10mM Tris-HCl pH8, 1mM EDTA). DNA was precipitated with 300mM of sodium acetate and 70% of chilled EtOH for 2 h at −20°C. DNA pellets were resuspended in formamide at 20ng/μL final concentration.

After performing classic immunofluorescence as described above (without the DAPI staining step), cells were post-fixed in 2% PFA for 12 minutes at RT°C and then permeabilized and deproteinized in PBS 0,5% Triton X-100 0,1M HCl for 10 minutes at RT°C. Samples were dehydrated in successive EtOH baths (2x 70% EtOH, 2x 85% EtOH and 2x 100% EtOH). After co-denaturation at 80°C for 5 minutes, cells’ DNA was hybridized with FISH probes diluted at 1/5 O/N at 37°C in dark and humid chamber. Cells were then washed 5 minutes in Saline-Sodium Citrate (SSC) 0,5X at 68°C, 2 minutes in SSC 1X at RT°C and incubated in DAPI diluted in SSC 2X for 5 minutes at RT°C. Coverslips were mounted in Fluoromount-G and stored at 4°C before observation.

### Microscopy, imaging, and quantification

Images were acquired with the Axio Observer Z1 inverted wide-field epifluorescence microscope (100X or 63X objectives/N.A. 1.46 or 1.4) (Zeiss) and a CoolSnap HQ2 camera from Photometrics. Identical settings and contrast were applied for all images of the same experiment to allow data comparison. Raw images were treated with Fiji software or with Photoshop (Adobe). HIRA complex accumulation in PML NBs was attested by manual counting of a minimum of 100 cells for each condition and per replicate. PML-NBs and genes loci proximity was measured using the Fiji RenyiEntropy mask on PML and FISH staining. X and Y coordinates for the center of the spots were recovered and all distances between each PML NBs and gene loci were calculated using the formula 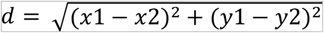 to find the minimal distance in each nucleus. Quantification of nuclear intensities was performed with Fiji. Briefly, DAPI and PML stainings were used to define masks of nuclei and of PML NBs. We quantified mean HA fluorescence intensity within each nucleus with the measure function applied on the red (HA) channel. To quantify HIRA intensity outside PML NBs, we first created a mask of nuclei devoid of PML NBs (Image calculator function of Fiji) and then applied the measure function on HIRA channel.

### Western blotting (WB)

Total cellular extracts were obtained by directly lysing the cells in 2X Laemmli sample buffer (LSB) (125 mM Tris-HCl pH 6.8, 20% glycerol, 4% SDS, bromophenol blue) containing 100mM DTT. RIPA extracts were obtained by lysing the cells in RIPA buffer (50mM Tris-HCl pH 7.5, 150mM NaCl, 0,5% Na-Deoxycholate, 1% NP-40, 0,1% SDS, 5mM EDTA) supplemented with 1X protease inhibitor cocktail (PIC) for 20min on ice. After incubation, RIPA extracts were centrifugated for 10min at 16000g at 4°C and supernatants were recovered and diluted with 4X LSB.

Western Blot was performed as in (Corpet et al. 2014) (see **Sup. Table 3** for antibodies dilution). Signal was revealed on ChemiDoc Imaging System (Bio-Rad) by using Amersham ECL Prime Western Blotting Detection Reagent (GE Healthcare Life Sciences, RPN2236) or Clarity Max Western ECL Blotting Substrate (Bio-Rad, 1705062).

### Chromatin immunoprecipitation (ChIP)

Cells were crosslinked directly in the culture dishes according to (Becker et al. 2017). After the PBS washes, cell pellets were snap-frozen in liquid nitrogen and stored at −80°C before immunoprecipitation. Cells were de-frozen on ice and chromatin was prepared following the TruChIP protocol from Covaris, as described in (Cohen et al. 2018). We used the Covaris M220 Focused-ultrasonicator to shear through chromatin (7 minutes at 140W, Duty off 10%, Burst cycles 200). After shearing, chromatin immunoprecipitation was performed as in (Becker et al. 2017). We used 20uL of protein A magnetic dynabeads (Invitrogen, 10001D) for immunoprecipitation with 2μg of the following rabbit primary antibodies: anti-H3.3 (Diagenode, C15210011), anti-panH3 (Abcam, ab1791), rabbit IgG (Diagenode, C15410206)). After DNA purification according to (Becker et al. 2017), DNA pellets were resuspended in ddH20 and stored at −20°C before qPCR analysis.

### Reverse Transcription (RT)

TRIzol reagent protocol (Invitrogen, 15596026) was used to isolate total RNAs, resuspended in ddH2O according to the manufacturer instructions. Contaminant DNA was removed with the DNA-free DNA Removal Kit (Invitrogen, AM1906). We used 1μg of RNA for reverse transcription (RT). RNAs were annealed with Random Primers (Promega, C118A) and RT was performed with the RevertAid H Minus Reverse Transcriptase (Thermo Scientific, EP0452) according to the manufacturer instructions. cDNAs were stored at −20°C before qPCR analysis.

### Quantitative PCR (qPCR)

qPCRs were performed using the KAPA SYBR qPCR Master Mix (SYBR Green I dye chemistry) (KAPA BIOSYSTEMS, KK4618). Primers used for qPCR are described in **Supplementary Table 4**.

### ChIP-Seq analysis

After ChIP, libraries were made in BGI and sequenced on a BGISEQ-500 sequencing platform (https://www.bgi.com). An average of 34 Million single-end 50bp reads was obtained for each library. Reads were trimmed using Trimmomatic and quality assessed with FastQC. Reads were aligned to the human genome hg38 using the BWA alignment software. Duplicate reads were identified using the picard tools script and only non-duplicate reads were retained. Broad peaks calling was performed with MACS2 (Zhang et al. 2008) (“--extsize 250 -q 0.01 --broad --broad-cutoff 0.05”), using input DNA as control. We defined all possible locations of H3.3 by merging broad peaks identified in our four conditions (n=190295), and annotated them with Homer (http://homer.ucsd.edu/homer/download.html). We counted reads extended to 250bp falling into these possible locations, in the four ChIP and their corresponding inputs, using bedtools-intersect. CPMs were obtained by dividing raw counts by the total number of mapped reads normalized to 1e6, and RPKMs by dividing CPMs by the peak length normalized to 1e3. Input RPKMs, used as background, was substracted from the respective ChIP RPKMs. We focused on 0,5% of the peaks with highest RPKM difference (n=951) between IFNβ treated and not treated conditions, of which 711 were intragenic. These peaks allowed us to defined a set of 654 genes, on which we performed GO analysis, with MsigDB, using enrichR plaform (Kuleshov et al. 2016).

As a complementary approach, we measured the ChIP enrichment within the 1000bp regions spanning the TESs (–500 +500), extending all unique reads into 250bp fragments, and counting those falling within TES using bedtools-intersect. CPMs were obtained similarly, and input DNA CPMs, used as background, was substracted from ChIP CPMs. Genes with the log2 of differential TES enrichment between IFNβ treated and not treated conditions being higher than 5 (log2(Fold Change)>5) were retained for GO analysis, as described above.

PlotProfile were generated using the DeepTools suite, starting from the MACS2 fold enrichment bigwig files, which take into account the read extension, the input DNA background and the library size normalization. The list of 48 core ISGs and 48 non-ISGs equal in size to the core ISGs was taken from (McFarlane et al. 2019). In order to reduce the noise on the profiles, we selected for each gene the transcript with the highest H3.3 enrichment at the TES in the IFNβ treated condition. Genome browser snapshots of H3.3 enrichment were generated using Integrative Genomics viewer (IGV : https://software.broadinstitute.org/software/igv/).

### Statistical analyses

Statistical analyses were performed using GraphPad Prism 6. To perform Student t test, we verified normal distribution of samples using Shapiro test and variance equality with Fisher test. Mann-Whitney u-test was applied in absence of normality for the sample distribution. p-values are depicted on graphs as follows: *<0,05; **<0,01; ***<0,001; ****<0,0001.

## Materials availability

All plasmids and cell lines generated in this study can be accessed upon request to the corresponding authors.

## Data availability

The ChIP-Seq datasets have been deposited in the Gene Expression Omnibus (GEO; http://www.ncbi.nlm.nig.gov/geo/) under the accession number GSE186937. The secure token to allow review of the GSE186937 record is: cjulkoaezbafpid

## Competing interest statetement

None declared.

## Acknowledgments

We thank Dr. Valérie Lallemand-Breitenbach for the *Pml* WT and *Pml−/−* MEFs. We thank Dr. Chris Boutell for the pLVX-His-SUMO1/2/3 plasmids. We thank Dr. Roger Everett for the PML-containing plasmids. We thank Dr. David J. Hughes for the SUMO-specific Affimers plasmids. We thank Lars Jansen for the pBABE-H3.3-SNAP-HA3 plasmid. We thank Dr. Caroline Schluth-Bolard for her kind help with the FISH on ISGs and for the GHRL BAC.

