## Supplementary information for "Interplay between PML NBs and HIRA for H3.3 dynamics following type I interferon stimulus"

#### SUPPLEMENTARY METHODS

**Supplementary Table 1: Lists of drugs and molecules used in this study**

| Name | Reference | Final concentration |
|---|---|---|
| Doxycycline | Sigma-Aldrich, D9891 | 100ng/mL |
| IFNβ (human, recombinant) | Peprotech, 300-02BC | 100 or 1000U/mL |
| IFNα (mouse, recombinant) | PBL assay science 12105-1 | 1000U/mL |
| IL-6 (human, recombinant) | Peprotech, 200-06 | 200ng/mL |
| IL-8 (human, recombinant) | Peprotech, 200-08 | 200ng/mL |
| PolyI:C | Invivogen, tlrl-pic | 10μg/mL |
| Ruxolitinib | Invivogen, tlrl-rux | 2μM |
| TNFα (human, recombinant) | Invivogen, rcyc-htnfa | 100ng/mL |
| Trichostatin A | Sigma T1952 | 2μM |

**Supplementary Table 2: List of siRNAs and their sequences**

| siRNA | Sequence | Reference |
|---|---|---|
| siHIRA | 5’GGAUAACACUGUCGUCAUCdTdT | (Ray-Gallet et al. 2011) |
| siLuc | 5’CGUACGCGGAAUACUUCGAdTdT | (Adam et al. 2013) |
| siPML | 5’AGATGCAGCTGTATCCAAGdTdT | (Everett et al. 2006) |
| siSP100 | 5'GUGAGCCUGUGAUCAAUAAdTdT | (Berscheminski et al. 2014) |
| siSUMO-1 | 5’GGACAGGAUAGCAGUGAGAdTdT | (Lallemand-Breitenbach et al. 2008) |
| siSUMO-2/3 | 5’GUCAAUGAGGCAGAUCAGAdTdT | (Yao et al. 2011) |

**Supplementary Table 3: List of primary antibodies**

| Antibody | Species | Reference | Dilution | |
|---|---|---|---|---|
| | | | IF | WB |
| Actin | Rabbit polyclonal | Sigma-Aldrich A2066 | | 1:1000 |
| panH3 | Rabbit polyclonal | Abcam ab1791 | | 1:5000 |

| | | | | |
|---|---|---|---|---|
| H3.3 | Rabbit monoclonal | Diagenode C15210011 | | 1:1000 |
| HA | Rabbit polyclonal | Abcam Ab9110 | 1:1000 | 1:1000 |
| HIRA #01 | Mouse monoclonal (clone WC119) | Active Motif 3558 | 1:500 | 1:1000 |
| HIRA #02 | Mouse monoclonal (clone WC119) | Millipore 04-1488 | 1:500 | 1:1000 |
| HIRA #03 (for mouse) | Rabbit polyclonal | Abcam ab20655 | 1:100 | |
| 6xHis #01 | Mouse monoclonal (clone 3D5) | Clontech 631212 | 1:1000 | |
| 6xHis #02 | Rabbit polyclonal | Bethyl A190-114A | 1:10000 | |
| c-Myc #01 | Mouse monoclonal (clone 9E10) | Santa Cruz sc-40 | | 1:1000 |
| c-Myc #02 | Rabbit polyclonal | Abcam ab9106 | 1:1000 | |
| PML #01 | Mouse monoclonal (clone PG-M3) | Santa Cruz sc-966 | 1:200 | |
| PML #02 | Rabbit polyclonal | Santa Cruz sc-5621 | 1:200 | 1:1000 |
| PML #03 | Rabbit polyclonal | Sigma PLA0172 | 1:5000 | 1:1000 |
| PML #04 (for mouse) | Mouse monoclonal | Millipore MAB3738 | 1:100 | |

| | | | | |
|---|---|---|---|---|
| | (clone 36.1-104) | | | |
| SUMO-1 | Rabbit monoclonal (clone Y299) | Abcam ab32058 | | 1:1000 |
| SUMO-2/3 | Rabbit polyclonal | Abcam ab3742 | | 1:1000 |
| αTubulin | Mouse monoclonal (clone DM1A) | Sigma T6199 | | 1:10000 |

**Supplementary Table 4: List of primers used for qPCR**

| **Name** | **Forward primers (5'→3')** | **Reverse primers (5'→3')** | **Ref** |
|---|---|---|---|
| **ChIP qPCR** | | | |
| H3.3-ChIP-cluster3-Chr1 (Enh1) | GCC-ACT-TGC-CAA-TGT-TTC-TC | TGG-CCC-CAT-GTA-GTG-AAA-AG | (Pchelintsev et al. 2013) |
| ChIP-MX1-TSS | GGG-ACA-GGC-ATC-AAC-AAA-GCC | GCC-CTC-TCT-TCT-TCC-AGG-CAA-C | (Cheon et al. 2013) |
| ChIP-MX1-mid | TCT-ACG-CTC-TGG-GGA-CAT-CA | GAA-CCA-AAC-CCA-CCA-CCA-GA | |
| ChIP-MX1-TES | CTC-CCG-TGA-ACT-GTT-CTT-TCC-T | GCT-GTA-GGT-GTC-CTT-GTC-CT | |
| ChIP-OAS1-TSS | ACC-ACA-GAC-AAC-TGT-GAA-AGG | GTC-CTT-TAG-CCA-GCA-ACA-AGC | |
| ChIP-OAS1-mid | GCA-GCA-CGT-TGG-GAG-ATA-GA | TTC-TCC-TGA-TGT-GGC-AAG-GG | |
| ChIP-OAS1-TES | CTT-GTC-ACA-TCC-CCA-CCT-CTC | GTC-CTT-TGC-CCC-TGT-TTA-GC | |
| ChIP-ISG54-TSS | GCA-GGA-AGT-GGG-GTT-TGC-TA | GAG-GGA-TGT-TTC-ATC-GGC-CT | |

| | | |
|---|---|---|
| ChIP-ISG54-mid | ATG-TAA-CTA-ACC-CCA-GGT-GCG | TGC-TTC-CCA-CTC-CCA-TTT-TGA |
| ChIP-ISG54-TES | AGT-CTG-GAA-GCC-TCA-TCC-CT | CCT-AGT-GGG-CAC-CAC-ATC-TC |
| **RT qPCR** | | |
| *GAPDH* | GAG-TCA-ACG-GAT-TTG-GTC-GT | TTG-ATT-TTG-GAG-GGA-TCT-CG |
| *H3F3A* | CCA-GGA-AGC-AAC-TGG-CTA-CA | ACC-AGG-CCT-GTA-ACG-ATG-AG |
| *HIRA* | AGG-ACT-CTC-GTC-TCA-TGC-CT | CAG-CTT-CAG-TGC-AAG-TGC-TG |
| *ISG15* | GGT-GGA-CAA-ATG-CGA-CGA-AC | TCG-AAG-GTC-AGC-CAG-AAC-AG |
| *ISG54* | TGA-AAG-AGC-GAA-GGT-GTG-CT | CTC-AGA-GGG-TCA-ATG-GCG-TT |
| *MX1* | GGA-GGC-ACT-GTC-AGG-AGT-TG | TCC-TGG-TAA-CTG-ACC-TTG-CC |
| *OAS1* | AGC-TGG-AAG-CCT-GTC-AAA-GA | AGG-TTT-ATA-GCC-GCC-AGT-CAA |
| *PML* | CAG-GGA-CCC-TAT-TGA-CGT-TG | ATG-GAG-AAG-GCG-TAC-ACT-GG |

#### Supplementary Figure 1

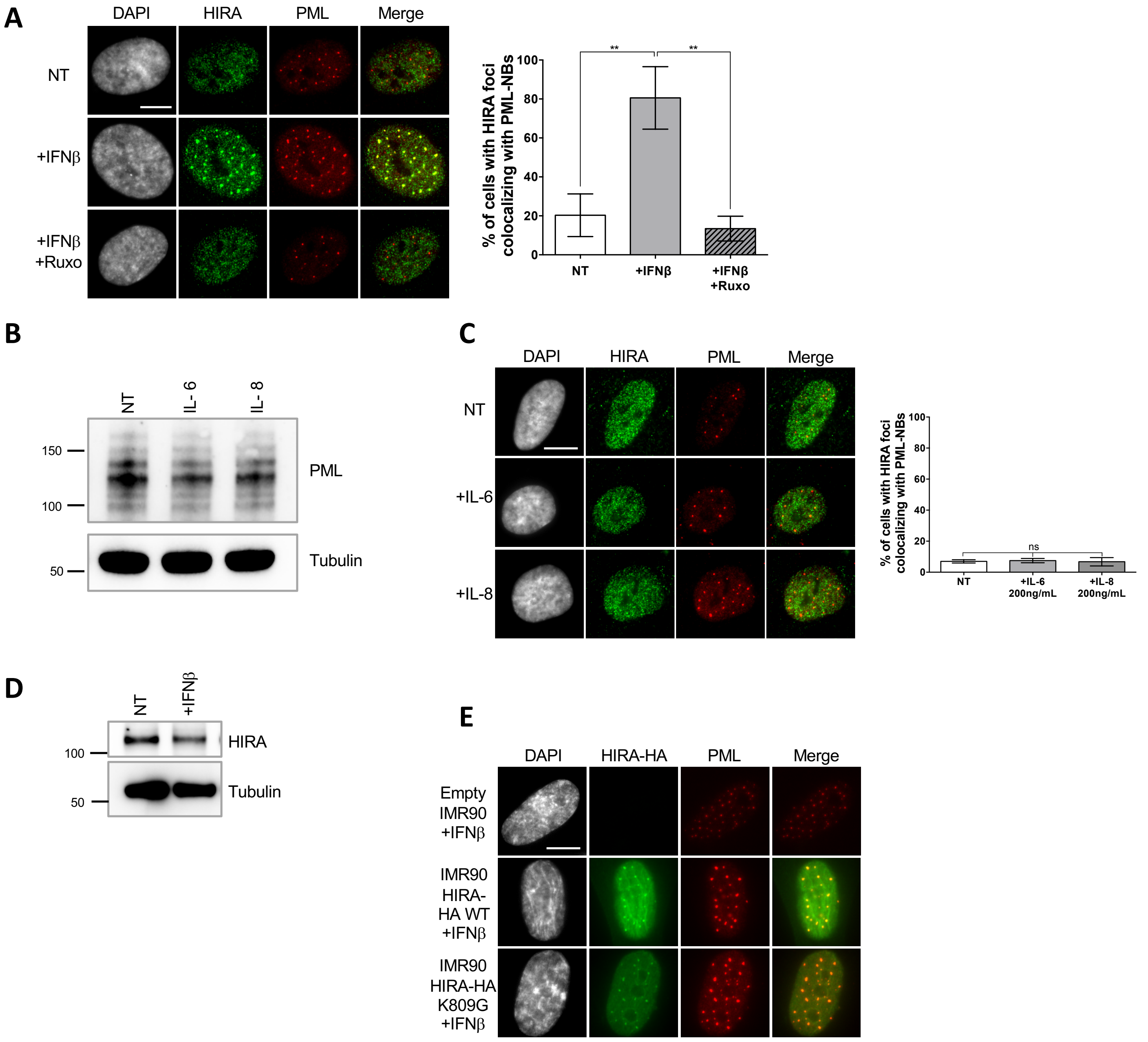

**Supplementary Figure 1. Impact of various cytokines on HIRA localization in PML NBs.**
**A.** (left) Fluorescence microscopy visualization (left) of HIRA (green) and PML (red) in BJ cells treated with IFNβ at 1000U/mL for 24h. Ruxolitinib (Ruxo) was added at 2μM one hour before IFNβ and left for 25h. Cell nuclei are visualized by DAPI staining (grey). Scale bar represents 10μm. (right) Histogram shows quantitative analysis of cells with HIRA localization at PML NBs. Numbers represent the mean of 3 independent experiments (±SD). p-values (Student t-test): **<0,01. **B.** Western blot visualization of PML in BJ cells treated with IL-6 or IL-8 at 200ng/mL for 24h. Tubulin is a loading control. **C.**(left) Fluorescence microscopy visualization (left) of HIRA (green) and PML (red) in BJ cells treated as in B. Cell nuclei are visualized by DAPI staining (grey). Scale bar represents 10μm. (right) Histogram shows quantitative analysis of cells with HIRA localization at PML NBs. Numbers represent the mean of 3 independent experiments (±SD). p-value (Student t-test): ns: non significant. **D.** Western blot visualization of HIRA in BJ cells treated as in A. Tubulin is a loading control. **E.** Fluorescence microscopy visualization of HA (green) and PML (red) in normal IMR90 cells ('empty') or in IMR90 cells stably expressing HIRA-HA WT or HIRA-HA mutated on K809G. Cells were treated with IFNβ at 1000U/mL for 24h. Cell nuclei are visualized by DAPI staining (grey). Scale bar represents 10μm.

#### Supplementary Figure 2

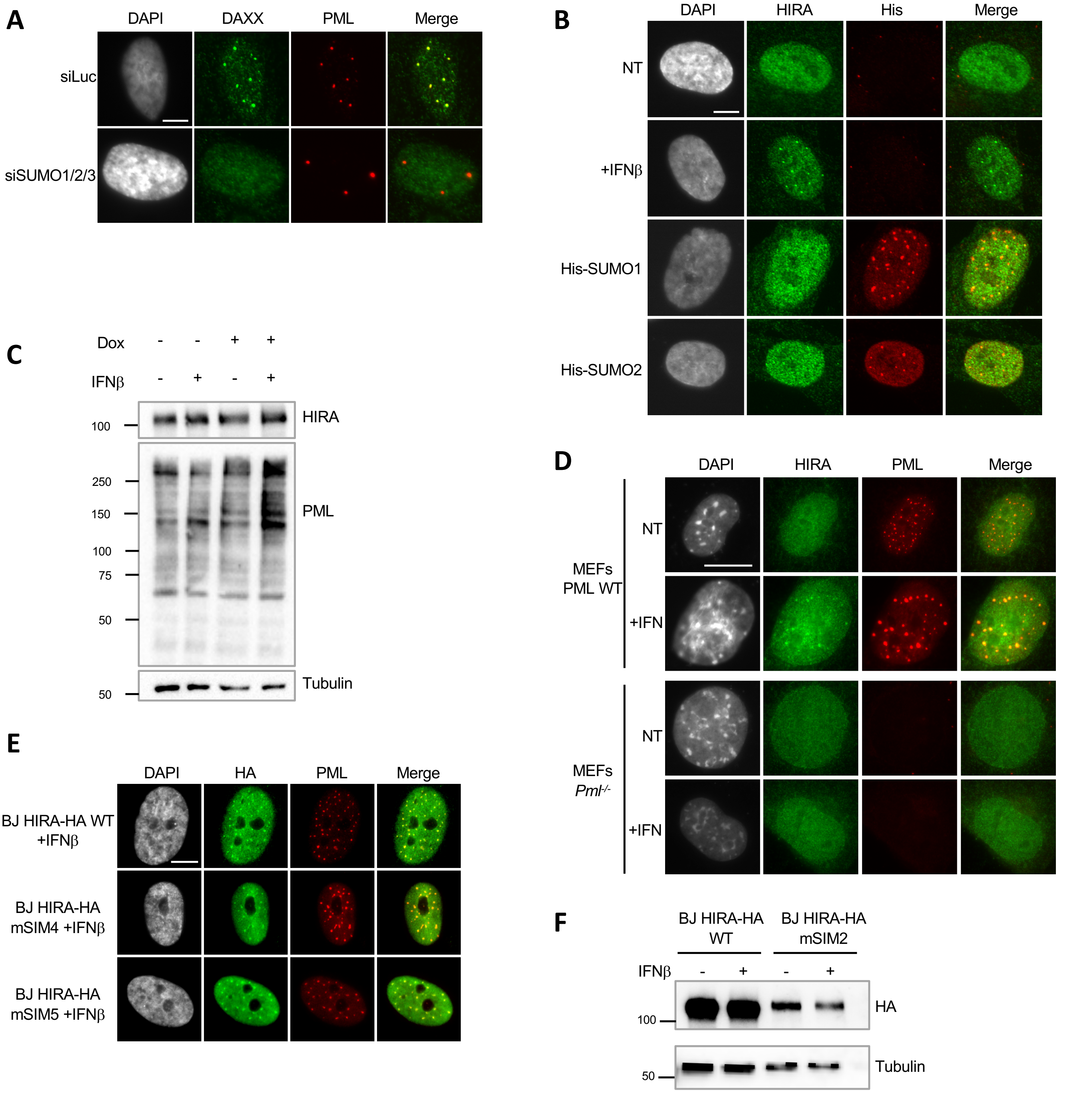

**Supplementary Figure 2. HIRA accumulation in PML NBs does not depend on overexpression of SUMO proteins but on a SIM motif.**

**A.** Fluorescence microscopy visualization of DAXX (green) and PML (red) in BJ cells treated with the indicated siRNAs for 48h. Cell nuclei are visualized by DAPI staining (grey). Scale bar represents 10mm. **B.** Fluorescence microscopy visualization of HIRA (green) and 6xHis-SUMO (red) in BJ cells transduced with lentiviral vectors 6xHis-tagged SUMO-1 or SUMO-2 proteins. Cells were fixed 48h after the transduction. Cell nuclei are visualized by DAPI staining (grey). Scale bar represents 10μm. **C.** Western blot visualization of HIRA and PML from total cellular extracts of S1S2D5-His Affimer-transduced BJ cells treated as in Figure 3C. Tubulin is a loading control. **D.** Fluorescence microscopy analysis of HIRA (green) and PML (red) in WT MEFs and in MEFs *Pml*$^{-/-}$. Cell nuclei are visualized by DAPI staining (grey). Scale bar represents 10μm. **E.** Fluorescence microscopy analysis of HIRA-HA (green) and PML (red) in BJ cells transduced with HIRA-HA WT, or mSIM4/5 mutants and treated with IFNβ for 24h. Cell nuclei are visualized in fluorescence microscopy by DAPI staining (grey). Scale bar represents 10μm. **F.** Western blot visualization of HA from total cellular extracts of BJ cells stably expressing HIRA-HA WT or HIRA-HA mSIM2. Tubulin is a loading control.

#### Supplementary Figure 3

**A**

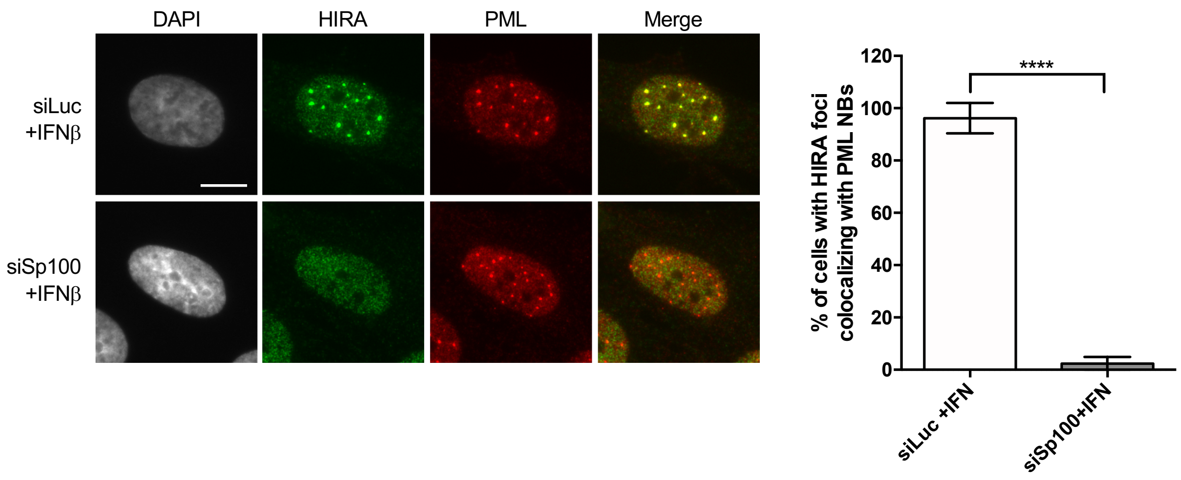

**B**

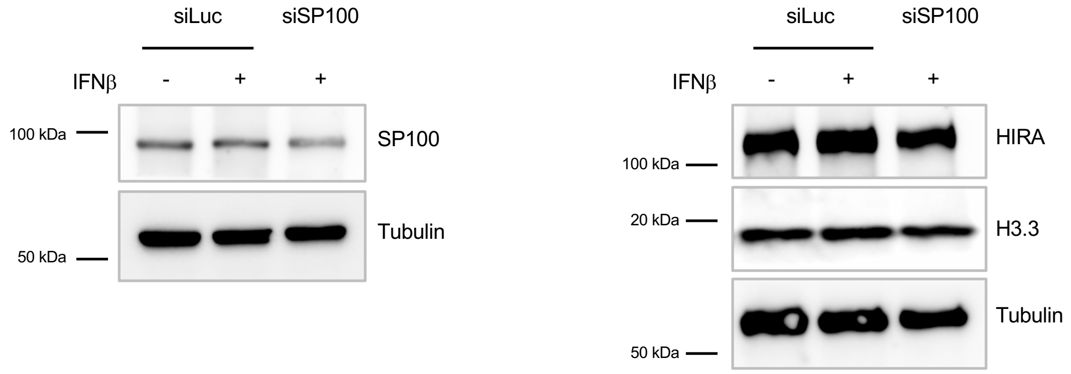

**C**

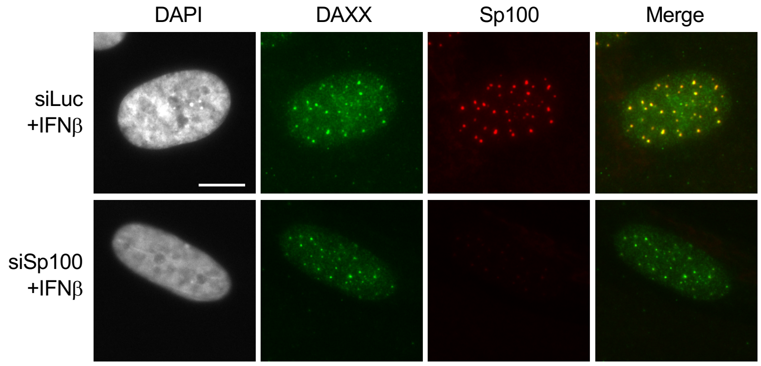

**Supplementary Figure 3. SP100 depletion impairs HIRA accumulation in PML NBs upon IFN-I treatment.**

**A.** (left) Fluorescence microscopy visualization of HIRA (green) and PML (red) in BJ cells treated with the indicated siRNAs for 48h and with IFNβ at 1000U/mL for the last 24h of siRNAs treatment. Cell nuclei are visualized by DAPI staining (grey). Scale bar represents 10mm. (right) Histogram shows quantitative analysis of cells with HIRA accumulation in PML NBs. Numbers represent the mean of 3 independent experiments (±SD). p-value (Student t-test): ****<0,0001. **B.** Western blot visualization of SP100, HIRA and H3.3 from total cellular extracts of BJ cells treated as in A. Tubulin is a loading control. **C.** Fluorescence microscopy visualization of DAXX (green) and Sp100 (red) in BJ cells treated with the indicated siRNAs for 72h and with IFNβ at 1000U/mL for the last 24h of siRNAs treatment. Cell nuclei are visualized by DAPI staining (grey). SP100 depletion does not impair the constitutive presence of DAXX in PML NBs. Scale bar represents 10mm.

### Supplementary Figure 4

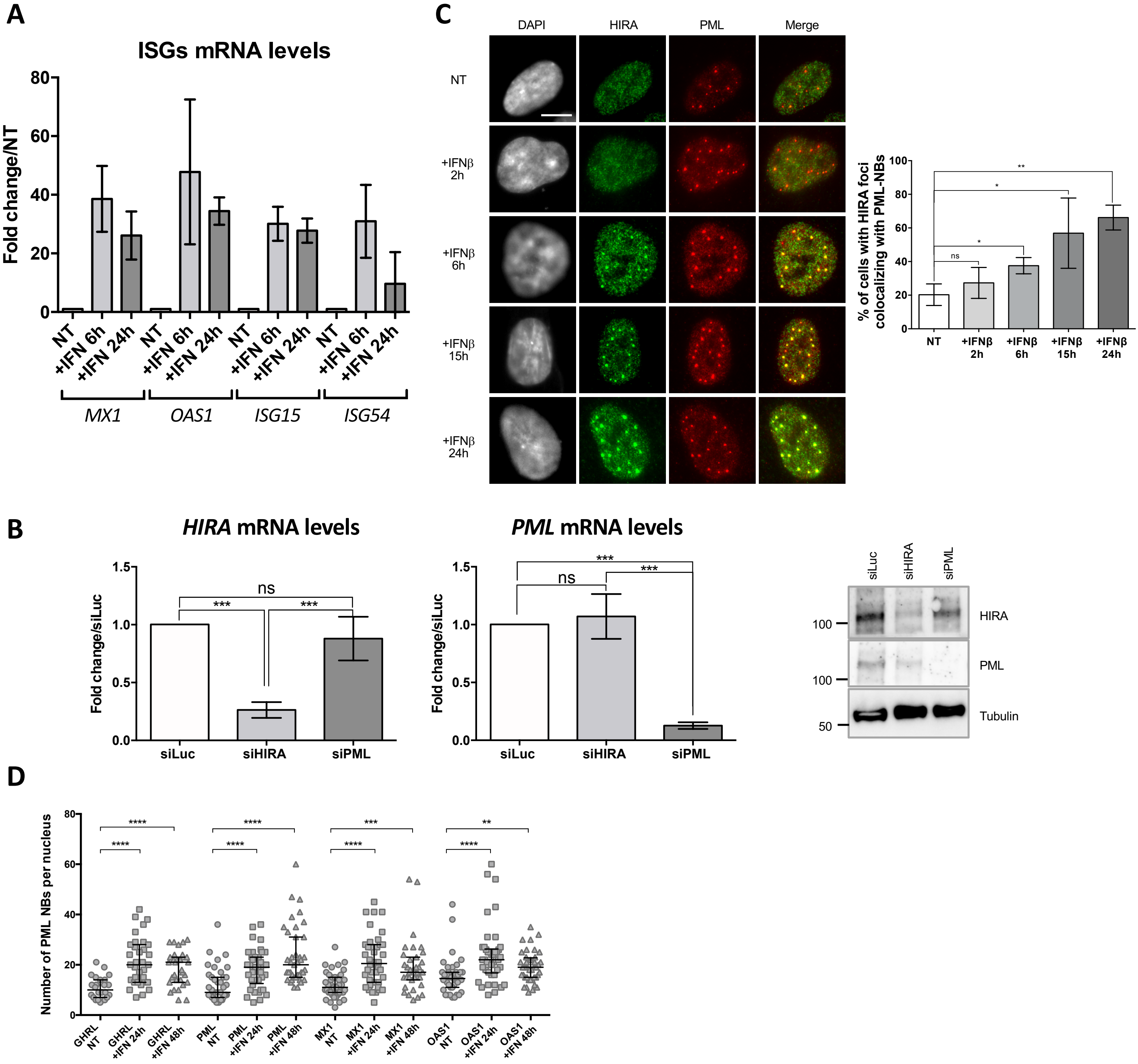

**Supplementary Figure 4. ISGs are upregulated upon IFN-I treatment and become juxtaposed to PML NBs.**

**A.** Histogram shows ISGs mRNA relative levels of *MX1*, *OAS1*, *ISG15* and *ISG54* normalized on *GAPDH* mRNA levels of BJ cells treated with IFNβ at 100U/mL for the indicated time. Rationalization was performed on mRNA levels of non-treated cells. Numbers represent the mean of 2 independent experiments. **B.** (left) Histograms show *HIRA* and *PML* mRNA relative levels normalized on *GAPDH* mRNA levels of BJ cells treated with the indicated siRNAs for 72h. Rationalization was performed on mRNA levels of siLuc cells. Numbers represent the mean of 3-4 independent experiments (±SD). p-values (Student t-test): ***<0,001; ns: non significant. (right) Western blot visualization of HIRA and PML from total cellular extracts of BJ cells treated with the indicated siRNAs for 72h. Tubulin is a loading control. **C.** (left) Fluorescence microscopy visualization of HIRA (green) and PML (red) in BJ cells treated with IFNβ at 1000U/mL for the indicated time. Cell nuclei are visualized by DAPI staining (grey). Scale bar represents 10μm. (right) Histogram shows quantitative analysis of cells with HIRA localization at PML NBs. Numbers represent the mean of 3 independent experiments (±SD). p-values (Student t-test): *<0,05; **<0,01; ns: non significant. **D.** Scatter plot shows the number of PML NBs per nucleus. The line in the middle represents the median of all observations. Nuclei used for the analysis are the same as described in Figure 4B.

### Supplementary Figure 5

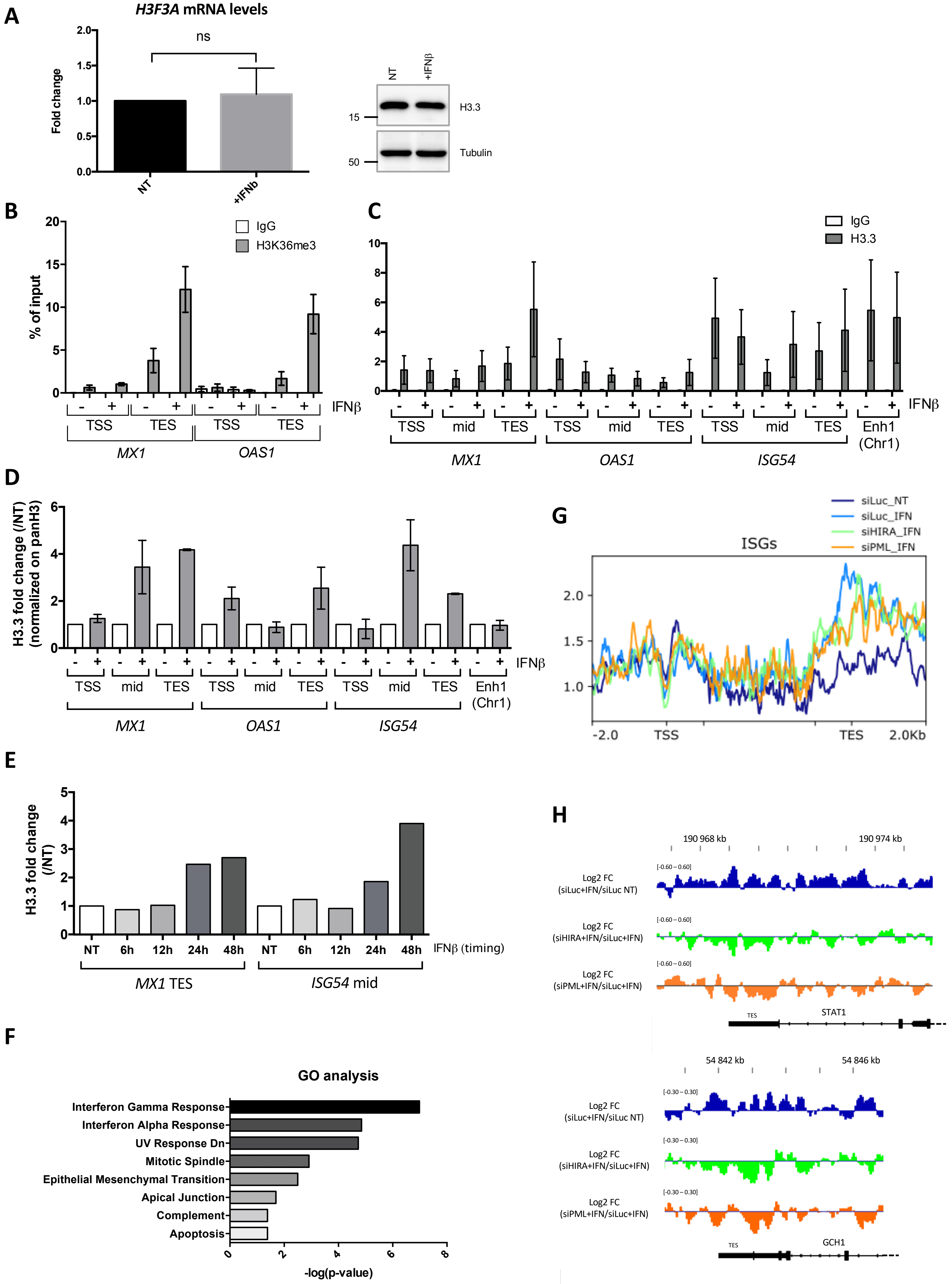

**Supplementary Figure 5. H3.3 and H3K36m3 increase on ISGs upon IFN-I.**
**A.** (left) Histogram shows relative *H3F3A* mRNA levels (normalized on *GAPDH* mRNA levels) of BJ cells treated or not with IFNβ at 1000U/mL. Rationalization was performed on the untreated condition. (right) Western blot visualization of H3.3 from total cellular extracts of BJ cells treated as in A. Tubulin is used here as a loading control. **B.** Histogram shows H3K36me3 and IgG enrichment (% of input) obtained through ChIP experiments on BJ cells treated or not with IFNβ at 1000U/mL for 24h. qPCR was performed on *MX1* and *OAS1* TSS and TES regions. Numbers represent the mean of 2 independent experiments (±SD). **C.** Histogram shows H3.3 and IgG enrichment (% of input) obtained through ChIP experiments on BJ cells treated or not with IFNβ at 1000U/mL for 24h. qPCR was performed on *MX1*, *OAS1* and *ISG54* ISGs TSS, mid and TES regions and on one enhancer region on chromosome 1 (Enh1). Numbers represent the mean of 3 independent experiments (±SD). **D.** Histogram shows H3.3 enrichment fold change obtained through ChIP experiments on BJ cells treated as in B. and normalized on panH3 enrichment. Rationalization was performed on H3.3 enrichment of untreated cells. qPCR was performed as in C. Numbers represent the mean of 2 independent experiments (±SD). **E**. Histogram shows H3.3 enrichment fold change obtained through ChIP experiments on BJ cells treated or not with IFNβ at 1000U/mL for the indicated times. Rationalization was performed on H3.3 enrichment in untreated cells. qPCR was performed on *MX1* TES and *ISG54* Mid. Numbers represent one experiment. **F.** Gene Ontology analysis on a set of 654 genes having the highest RPKM difference between IFNb treated and not treated conditions (see Materials and Methods). **G.** ChIP-Seq profile of H3.3 enrichment over 49 core ISGs (McFarlane et al. 2019) ranging from -2kb before TSS to 2kb downstream the TES in BJ cells treated as in Figure 5E. **H.** Representative genome browser snapshots of the H3.3 enrichment across the TES region of 2 ISGS : *STAT1* and GTP cyclohydrolase 1 (*GCH1*). Shown are the log2 fold changes of H3.3 enrichment in siLuc + IFN/siLuc NT cells (blue), siHIRA+IFN/siLuc+IFN treated cells (green) and siPML+IFN/siLuc+IFN treated cells (orange).

### Supplementary Figure 6

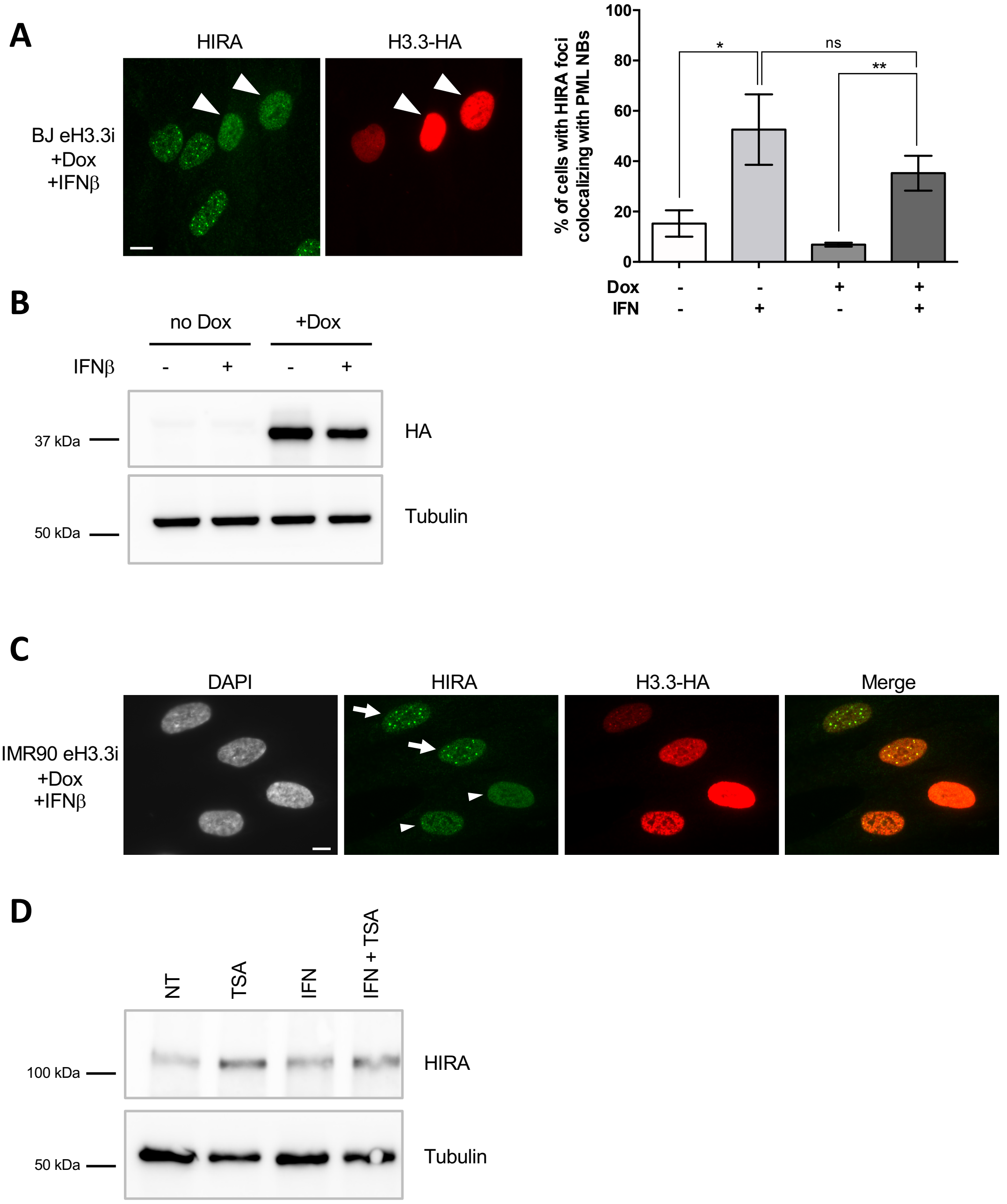

**Supplementary Figure 6. HIRA accumulation in PML NBs upon IFN-I treatment can be modulated by the pool of soluble H3.3-H4 and by chromatin compaction.**
**A.** (left) Fluorescence microscopy visualization of HIRA (green) and H3.3-HA (red) in BJ eH3.3i cells treated with doxycyclin (Dox) and IFNβ at 1000U/mL for 24h. Arrowheads indicate nuclei with high levels of nucleoplasmic H3.3-HA preventing HIRA accumulation in PML NBs despite IFNβ treatment. (right) Histogram shows quantitative analysis of cells with HIRA localization at PML NBs in BJ eH3.3i treated or not with Dox and IFNβ for 24h. p-values (Student t-test): *<0,05; **<0,01; ns: non significant. Numbers represent the mean of 3 independent experiments (±SD). **B.** Western blot visualization of HA from total cellular extracts of BJ eH3.3i cells treated as in A. Tubulin is a loading control. **C.** Fluorescence microscopy visualization of HIRA (green) and H3.3-HA (red) in IMR90 eH3.3i treated with Dox and IFNβ at 1000U/mL for 24h. Arrows indicate nuclei showing accumulation of HIRA in PML NBs together with H3.3-HA, while arrowheads indicate nuclei with high levels of nucleoplasmic H3.3-HA preventing HIRA accumulation in PML NBs despite IFNβ treatment. **D.** Western blot visualization of HIRA from total cellular extracts of BJ cells treated with TSA at 2μM 1h before IFNβ at 1000U/mL for 24h. Tubulin is used here as a loading control. **E.** Fluorescence microscopy visualization of HIRA-HA (green) and PML (red) in HIRA-HA WT/S697A transduced BJ cells treated with IFNβ at 1000U/mL for 24h. **A, C, E.** Cell nuclei are visualized by DAPI staining (grey). Scale bars represent 10μm.
